# Adult medial habenula neurons require GDNF receptor GFRα1 for synaptic stability and function

**DOI:** 10.1101/2021.07.07.451421

**Authors:** Diana Fernández-Suárez, Favio A. Krapacher, Katarzyna Pietrajtis, Annika Andersson, Lilian Kisiswa, Marco A. Diana, Carlos F. Ibáñez

## Abstract

The medial habenula (mHb) is an understudied small brain nucleus linking forebrain and midbrain structures controlling anxiety and fear behaviors. The mechanisms that maintain the structural and functional integrity of mHb neurons and their synapses remain unknown. Using spatio-temporally controlled Cre-mediated recombination in adult mice, we found that the GDNF receptor alpha 1 (GFRα1) is required in adult mHb neurons for synaptic stability and function. mHb neurons express some of the highest levels of GFRα1 in the mouse brain, and acute ablation of GFRα1 results in loss of septo-habenular and habenulo-interpeduncular glutamatergic synapses, with the remaining synapses displaying reduced numbers of presynaptic vesicles. Chemo- and opto-genetic studies in mice lacking GFRα1 revealed impaired circuit connectivity, reduced AMPA receptor postsynaptic currents, and abnormally low rectification index of AMPARs, suggesting reduced Ca^2+^-permeability. Further biochemical and proximity ligation assay studies defined the presence of GluA1/GluA2 (Ca^2+^-impermeable) as well as GluA1/GluA4 (Ca^2+^-permeable) AMPAR complexes in mHb neurons, as well as clear differences in the levels and association of AMPAR subunits in mHb neurons lacking GFRα1. Finally, acute loss of GFRα1 in adult mHb neurons reduced anxiety-like behavior and potentiated context-based fear responses, phenocopying the effects of lesions to septal projections to the mHb. These results uncover an unexpected function for GFRα1 in the maintenance and function of adult glutamatergic synapses, and reveal a potential new mechanism for regulating synaptic plasticity in the septo-habenulo-interpeduncular pathway and attuning of anxiety and fear behaviors.

## Introduction

The habenula is a phylogenetically conserved brain structure, present in all vertebrates, consisting of two small nuclei located above the thalamus, close to the midline. Due to its distinct location and connectivity, it is thought to function as a node linking more recently evolved forebrain structures involved in executive functions with ancient midbrain areas that process aversion and reward (1). Lesions to the habenula affect different behaviors related to stress, fear, aversion, anxiety, pain and sleep (2–5). In mammals, the habenula is subdivided into two distinct subnuclei, the lateral (LHb) and medial (mHb) habenula, that present different neurochemical properties, connectivity and functions (1). In the fasciculus retroflexus (FR), one of the longest major fiber tracts in the brain and among the first to form during development, afferent axons from the two subnuclei are segregated, with LHb axons in the periphery, innervating different midbrain structures, and mHb axons in the center, all terminating in the interpeduncular nucleus (IPN) (1). Perhaps due to its prominent connections to dopaminergic and serotoninergic neurons in ventral tegmental area (VTA), substantia nigra pars compacta (SNpc) and raphe nuclei, the LHb has been more extensively studied than the mHb (6). However, mHb neurons display several characteristics that make them quite unique in the mammalian brain. A striking property of mHb neurons is their spontaneous pacemaking activity, which generates tonic trains of action potentials of about 2-10 Hz (7). Moreover, GABAergic inputs from the medial septum elicit excitation, rather than inhibition, in mHb neurons due to lack of the KCl co-transporter KCC2, a feature that is more common in the developing brain, but rare among adult CNS neurons (8). Similarly, GABA released by postsynaptic IPN neurons has been shown to act as a retrograde messenger on presynaptic GABA-B receptors in mHb terminals to amplify glutamatergic transmission at habenulo-interpeduncular synapses (5,9). Furthermore, the mHb is the only known structure in the adult brain that contains functional NMDA receptor GluN1/GluN3A, through which glycine, otherwise known as a major inhibitory neurotransmitter, regulates neuronal excitability in mHb neurons (10). Finally, while the majority of AMPA receptors in the adult brain are Ca^2+^ impermeable, inputs to the mHb from the triangular septum (TS) and bed nucleus of the anterior commissure (BAC), as well as the mHb output to the IPN, produce glutamatergic responses through Ca^2+^ permeable AMPA receptors (9,11). Despite its unique properties and functional importance in fear and anxiety, very little is known about the mechanisms that regulate the structural and functional plasticity of mHb neurons and their synapses. Moreover, aside from electrophysiological evidence, the molecular composition and functional importance of mHb Ca^2+^-permeable AMPA receptors remain to be established.

GFRα1 is the main binding receptor for GDNF (glial cell line-derived neurotrophic factor). It lacks an intracellular domain and is anchored to the plasma membrane through a glycosyl-phosphatidylinositol link (12). Through the action of plasma membrane lipases, glycosyl-phosphatidy cleavage results in receptor shedding from the cell surface, allowing GFRα1 to act on nearby cells (13,14). GFRα1 has been shown to function both in conjunction with transmembrane co-receptor subunits, such as the receptor tyrosine kinase RET (12) and the neural cell adhesion molecule NCAM (15), as well as independently of these receptors (16). During development, GDNF and GFRα1 are essential molecular determinants for neuronal survival, migration and differentiation of a variety of neuronal subpopulations in the peripheral and central nervous systems (17). Exogenously provided GDNF can promote survival of midbrain dopaminergic neurons, both *in vitro* and in animal models of Parkinson’s Disease, and therefore has been tested in clinical trials for that disease (18). On the other hand, endogenous GDNF appears to be dispensable for the survival of adult dopaminergic neurons in mice (19). Although GFRα1 is highly expressed in several areas of the adult brain, such as the septum, midbrain and habenula (20–23), its function in the adult nervous system remains unknown.

The prominent expression of GFRα1 in the adult mHb led us to investigate its function in these neurons using temporally and spatially controlled Cre-mediated inactivation of the *Gfra1* gene. In this study, we report a previously unknown and essential role for GFRα1 in the maintenance, integrity and function of mHb synapses in the adult septum→mHb→IPN pathway. Significantly, loss of GFRα1 in the mHb resulted in changes in the molecular composition of mHb AMPA receptors that reduced their Ca^2+^ permeability, as well as alterations in fear and anxiety behaviors that paralleled those observed after lesion of the septo-habenular pathway (4).

## Results

### Characterization of GFRα1 and co-receptor expression in the septum→Hb→IPN pathway

Inspection of sections through the brain of tamoxifen-injected 3 month old mice carrying a *Rosa26*^dTOM^ reporter under the control of a *Gfra1*^Cre-ERT2^ allele (herein referred to as *Gfra1*^dTOM^) revealed very high dTOM signal in the mHb and the FR (Fig. 1A), comparable to or exceeding that observed in better known sites of GFRα1 expression, such as septum, SNpc or VTA (Fig, S1A). Particularly striking was the intensity of dTOM signal in mHb axons of the FR and their terminals in the IPN (Figs. 1A and S1A). Q-PCR analysis of *Gfra1* mRNA expression in regions microdissected from adult mouse brain showed highest expression in the Hb (Fig. S1B), while Western blots confirmed highest expression in the IPN (Fig. S1C), most likely derived from mHb axon terminals, since *Gfra1* mRNA was at background levels in this structure (Fig. S1B). Expression of *Gdnf* mRNA and GDNF protein was found across a number of adult brain regions, including Hb, septum and IPN (Figs. S1D, E). As GDNF is a soluble, diffusible ligand, it is likely to be available to most neurons in those structures.

**Fig 1.**
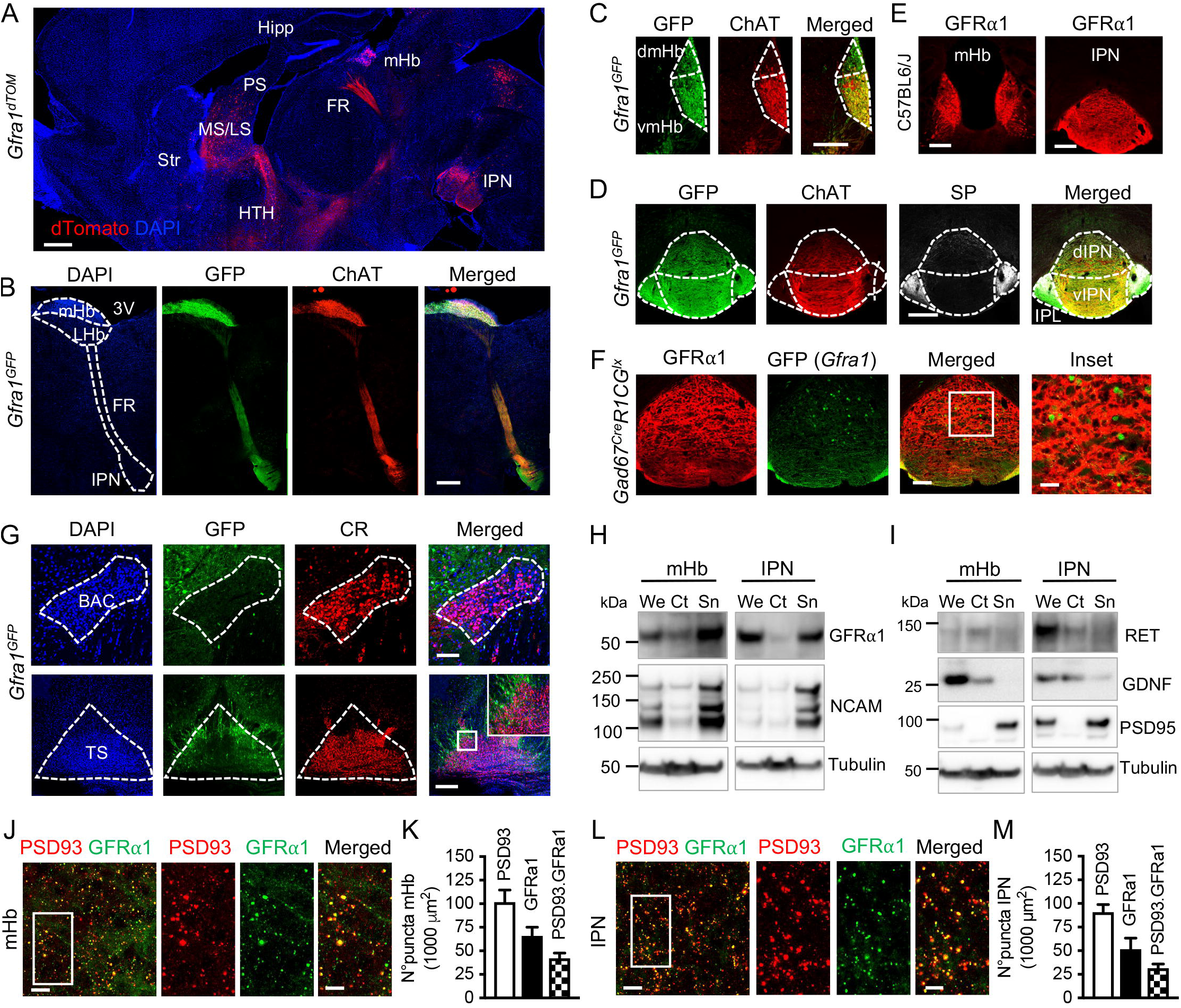
Characterization of GFRα1 and co-receptor expression in neurons and synapses of the septum→Hb→IPN pathway. A) dTomato epifluorescence (red) in sagittal sections of *Gfra1^dTOM^* mouse brain injected with tamoxifen at 3 months counterstained with DAPI (blue). FR: fasciculus retroflexus; Hipp: hippocampus; HTH: hypothalamus; IPN: interpeduncular nucleus; LS: lateral septum; mHb: medial habenula; MS; medial septum; Str: striatum. Scale bar, 500μm. B) GFP (green) and ChAT (red) immunostaining in sagittal sections of 3 month old *Gfra1^GFP^* mouse brain counterstained with DAPI (blue). 3V: 3th ventricle; LHb: lateral habenula. Scale bar, 400μm. C-D) GFP (green), ChAT (red) and SP (grey) immunostaining in coronal sections of the mHb (C) and IPN (D) of 3 month old *Gfra1^GFP^* mouse brain. dmHb: dorsal medial habenula; vmHb: ventral medial habenula: dIPN: dorsal IPN, vIPN: ventral IPN; IPL: lateral IPN. Scale bars, 200μm. E) GFRα1 immunostaining (red) in coronal sections of the mHb and IPN from 3 month old control (C57BL6/J) mouse. Scale bars, 200μm. F) GFP (green) and GFRα1 (red) immunostaining in coronal sections through the IPN of a 3 month old *Gad67^Cre^;R1CG^fx/fx^* mouse. Scale bars, 100μm (merged) and 35μm (inset). G) GFP (green) and calretinin (CR, red) immunostaining in coronal sections of the BAC and TS from 3 month old *Gfra1^GFP^* mouse brain counterstained with DAPI (blue). Scale bars, 100μm (BAC) and 200μm (TS). H,I) Immunoblots of whole (We), cytosolic (Ct) and synaptosome (Sn) protein extracts from the mHb and IPN of 3 month old C57BL6/J mice probed for GFRα1 and NCAM (H) or RET, GDNF and PSD95 (I). Tubulin was probed as loading control. J, L) GFRα1 (green) and PSD93 (red) immunostaining in the mHb (J) and IPN (L) of 3 month old C57BL6/J mice. Scale bars, 10μm (merged) and 5μm (inset). K, M) Quantification (±SEM) of GFRα1, PSD93 and double-labeled puncta in mHb (K) and IPN (M). N=5 mice (25-30 images per mouse).

For a closer inspection of the cellular distribution of the expression of GFRα1 and its co-receptors in the adult Hb, IPN and septum, we used mouse strains expressing GFP from either the *Gfra1* (*Gfra1*^GFP^) (see Methods section) or *RET* (*RET*^GFP^) (24) loci, in addition to *Gfra1*^dTOM^ mice and immunohistochemistry for endogenous GFRα1 and NCAM. All neurons in the mHb were positive for *Gfra1*^GFP^, while no GFP signal could be detected in the LHb (Figs. 1B, C). *Gfra1*^GFP^ signal co-localized with choline acetyltransferase (ChAT) in the ventral mHb (vmHb) (Figs. 1B, C). As it is well known, substance P (SP) can be detected at very low levels in dorsal mHb (dmHb) somas due to its rapid transport to axon terminal (25). In the IPN, both SP^+^ fibers, from the dmHb, as well as ChAT^+^ fibers, from the vmHb, displayed strong GFP signal (Fig. 1D). GFRα1 immunohistochemistry confirmed strong expression in the somas of adult mHb neurons and their axons throughout the IPN (Fig. 1E). To assess GFRα1 expression in the GABAergic IPN neurons, we used *R1CG* mice, expressing GFP from the *Gfra1* locus upon Cre-mediated recombination (26), bred to *Gad67*^Cre^ mice, which express Cre in GABAergic neurons (27). This revealed a small and sparse population of GFP^+^ cells in the dorsal IPN which did not appear labeled for GFRα1 immunohistochemistry (Fig. 1F), possibly reflecting GFP that persisted after transient expression of GFRα1 during development. At any rate, absence of a significant population of GFRα1 expressing cells in the IPN is in agreement with undetectable levels of *Gfra1* mRNA in this structure (Figs. S1B, see also S3H). As for GFRα1 co-receptors, NCAM was abundant in both the LHb and mHb subnuclei, overlapping with the *Gfra1*^dTOM^ signal in the latter (Fig. S1F). In contrast, *RET*^GFP^ signal was only detected in the LHb, but not in the mHb (Fig. S1F). Axonal projections labeled by *Gfra1*^dTOM^ and *RET*^GFP^ also appeared segregated in the FR, in agreement with the peripheral localization of LHb axons (28) (Fig. S1F). NCAM was detected throughout the IPN, while *RET*^GFP^ signal was only observed in scattered cells localized to the dorsal portion without significant overlap with mHb axons labeled by *Gfra1*^dTOM^ (Fig. S1G). In the septum, no *Gfra1*^GFP^ signal could be detected in calretinin (CR)^+^ neurons of the BAC or TS, the two subpopulations that project to the mHb (Fig. 1G). To ascertain whether the remaining *Gfra1*^GFP^ cells observed adjacent to the TS projected to the mHb, we performed flourogold retrograde tracing by stereotaxic injection in the mHb, followed by examination of fluorogold co-localization with *Gfra1*^GFP^ signal in the TS (Figs. S2A, B). Fluorogold did not overlap with GFP in the TS, although it did with CR, as expected (Figs. S2C, D), indicating that septal axons projecting to the mHb do not carry GFRα1. In the TS, a sparse cell subpopulation displayed *RET*^GFP^ signal, but it did not overlap with either the fluorogold tracer injected in the mHb (Figs. S2E-H), *Gfra1*^dTOM^ signal (Fig. S2I), or CR (Figs. S2G,H). No *RET*^GFP^ signal could be detected in BAC (Fig. S2G). Finally, NCAM was readily detected in both TS and BAC, where it decorated the plasma membrane of CR^+^ projection neurons (Figs. S2J, K).

Next, we assessed whether GFRα1 is located at synapses in the septum→mHb→IPN pathway. By Western blotting, we established the enrichment of GFRα1 and NCAM in synaptosomal fractions purified from adult mHb and IPN whole tissue extracts, as identified by the synaptic marker PSD95 (Figs. 1H, I). RET could not be detected in mHb extracts, and was excluded from synaptosomes in the IPN, while GDNF was present in whole tissue extract and cytosolic fractions of both structures (Fig. 1I). By immunohistochemistry, approximately two thirds of all GFRα1^+^ puncta in the mHb and in the IPN co-localized with the synaptic marker PSD93 (Figs. 1J-M), confirming a predominantly synaptic location of GFRα1 in these structures. Although the resolution of this analysis does not allow to determine the sub-synaptic localization of GFRα1, the absence of GFRα1 from septal neurons projecting to the mHb, and from IPN neurons, would indicate that GFRα1 is postsynaptic in septo-habenular synapses and presynaptic in habenulo-interpeduncular synapses.

### Adult ablation of GFRα1 induces presynaptic alterations and loss of glutamatergic synapses in the mHb and IPN

Global ablation of GFRα1 expression in adult mice was attained by crossing *Gfra1*^fx/+^ and *Gfra1*^CreERT2/+^ strains to obtain *Gfra1*^+/+^, *Gfra1*^CreERT2/+^ and *Gfra1*^CreERT2/fx^ animals, herein referred to as WT, Het and KO, respectively. Tamoxifen treatment at 3 months of age reduced GFRα1 expression to near background levels in the mHb and IPN of KO mice (Figs. S3A-H). Specific ablation of GFRα1 expression in the mHb was achieved by bilateral, stereotaxic injection of adeno-associated viruses (AAV2) expressing CRE recombinase under the control of a CMV promoter into the mHb of 3 month old *Gfra1*^fx/fx^ mice (herein called mHb.KO). Virus injection reduced GFRα1 levels by 56% and 60% in mHb and IPN, respectively, compared to mock-injected animals (herein called mHb.WT) (Figs. S3I-K).

The numbers of synaptic puncta double-labeled by pre- and post-synaptic markers Synapsin 1 and PSD93, respectively, were decreased by approximately half in both mHb and IPN of KO mice compared to Het or WT 28 days after tamoxifen treatment (Figs. 2A, B), suggesting loss of synapses after global ablation of GFRα1. A very pronounced decrease in the number of glutamatergic puncta labeled by vesicular glutamate transporters 1 and 2 (VGlut1 and VGlut2) was also observed in the mHb and IPN of KO mice 28 days after tamoxifen administration (Figs. 2C, D and S4A, B). In contrast, mHb synaptic puncta labeled by vesicular GABA transporter (VGAT) and gephyrin, corresponding to inhibitory synapses, were not affected (Fig. S4C, D). This indicated a specific loss of glutamatergic synapses in the KO mHb, which constitute the main afferents from the posterior septum (11,29). Importantly, the numbers of synaptic puncta labeled by Synapsin 1, PSD93, VGlut1 and VGlut2 were all significantly reduced in the mHb and IPN of mHb.KO mice 28 days after Cre virus injection (Figs. 2E-H), confirming the specific requirement of adult mHb GFRα1 expression for the maintenance of glutamatergic synapses in both the mHb and IPN.

**Fig 2.**
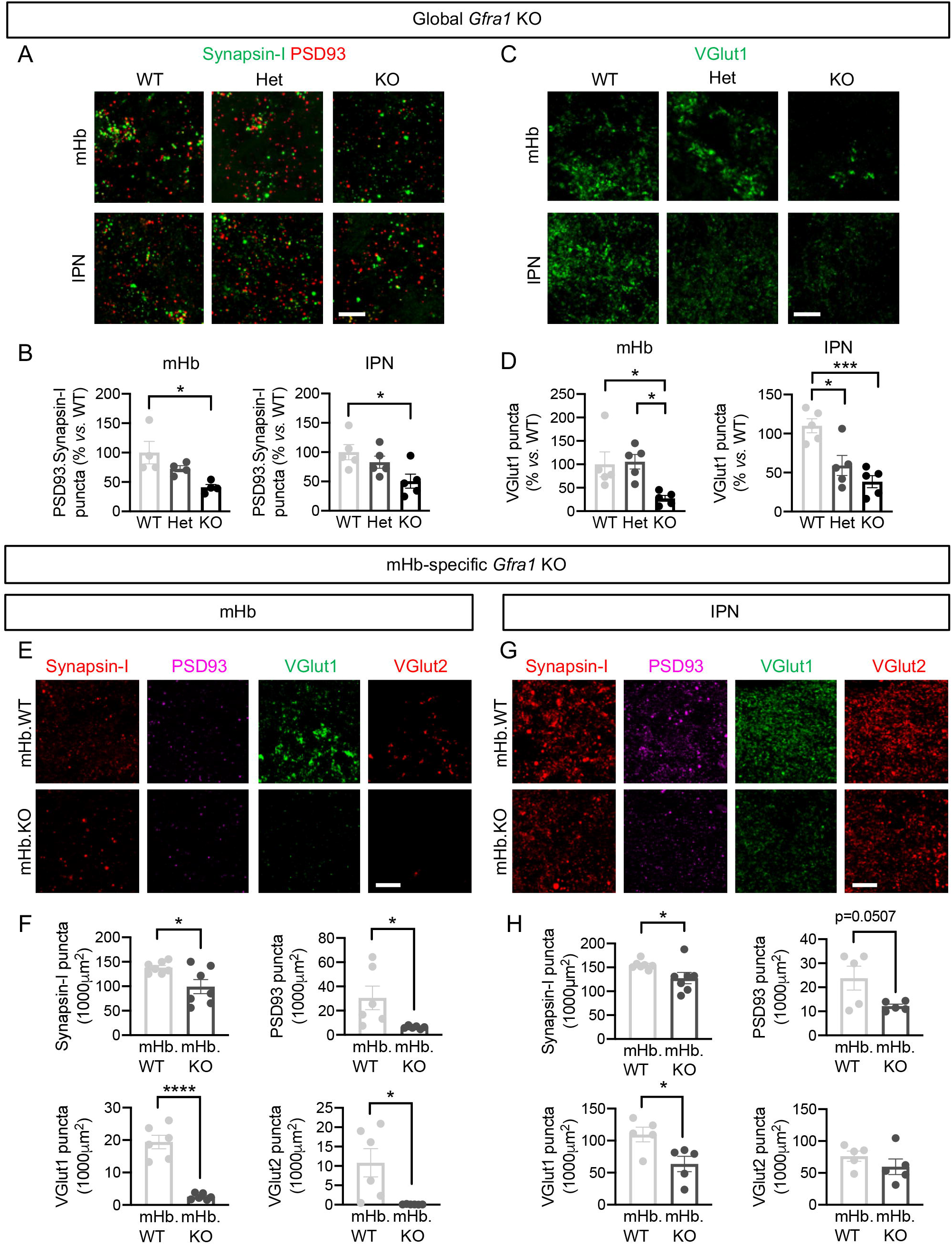
Global and mHb-specific ablation of GFRα1 induces loss of glutamatergic synapses in mHb and IPN neurons. A) Synapsin-I (green) and PSD93 (red) immunostaining in coronal sections of the mHb and IPN of WT, Het and KO mice. Scale bar, 10μm. B) Quantification (±SEM) of puncta with co-localized immunoreactivity for Synapsin-I and PSD93 in mHb and IPN. N=4-5 mice per group (25-30 images per mouse); 1-way ANOVA followed by Tukey’s post-hoc test; *, P <0.05. C) VGlut1 (green) immunostaining in coronal sections of the mHb and IPN of WT, Het and KO mice. Scale bar, 10μm. D) Quantification (±SEM) of VGlut1 puncta in mHb (K) and IPN in mHb and IPN. N=5 mice per group (25-30 images per mouse); 1-way ANOVA followed by Tukey’s post-hoc test; *, P <0.05; ***, P<0.001. E, G) Synapsin-I (red), PSD93 (purple), Vglut1 (green) and Vglut2 (red) immunostaining in coronal sections of the mHb (E) and IPN (G) in mHb.KO or mHb.WT mice. Scale bar, 10 μm. F, H) Quantification (±SEM) of immunoreactive puncta for each marker in mHb (E) and IPN (G) of mHb.KO or mHb.WT mice. N=6-7 mice per group (25-30 images per mouse); Student’s T-test; *, P< 0.05; ****, P< 0.0001.

Synapses in the mHb and IPN were further characterized by transmission electron microscopy. In agreement with our immunohistochemistry results, the number of synapses in the mHb was decreased by half in KO mice compared to Het or WT animals (Fig. 3A, B). Intriguingly, a marked presynaptic phenotype was observed in the remaining synapses, characterized by a reduction in the number of presynaptic vesicles, which affected both KO and Het mice (Fig. 3A, B). On the other hand, the postsynaptic length and the width of the synaptic cleft were not affected (Fig. 3A, B). Similar defects were observed in S and crest synapses in the IPN, previously shown to correspond to mHb afferents (30) (Fig. 3C, D). We note that there were no differences in mHb neuron number between WT, Het and KO mice 4 weeks after tamoxifen injection (Figs. S4E, F), indicating no effects on mHb neuron survival after acute ablation of GFRα1 in adult mice. Together, these data indicate that adult expression of GFRα1 in the mHb is essential for the maintenance of both the number and structure of glutamatergic synapses in the mHb and IPN.

**Fig 3.**
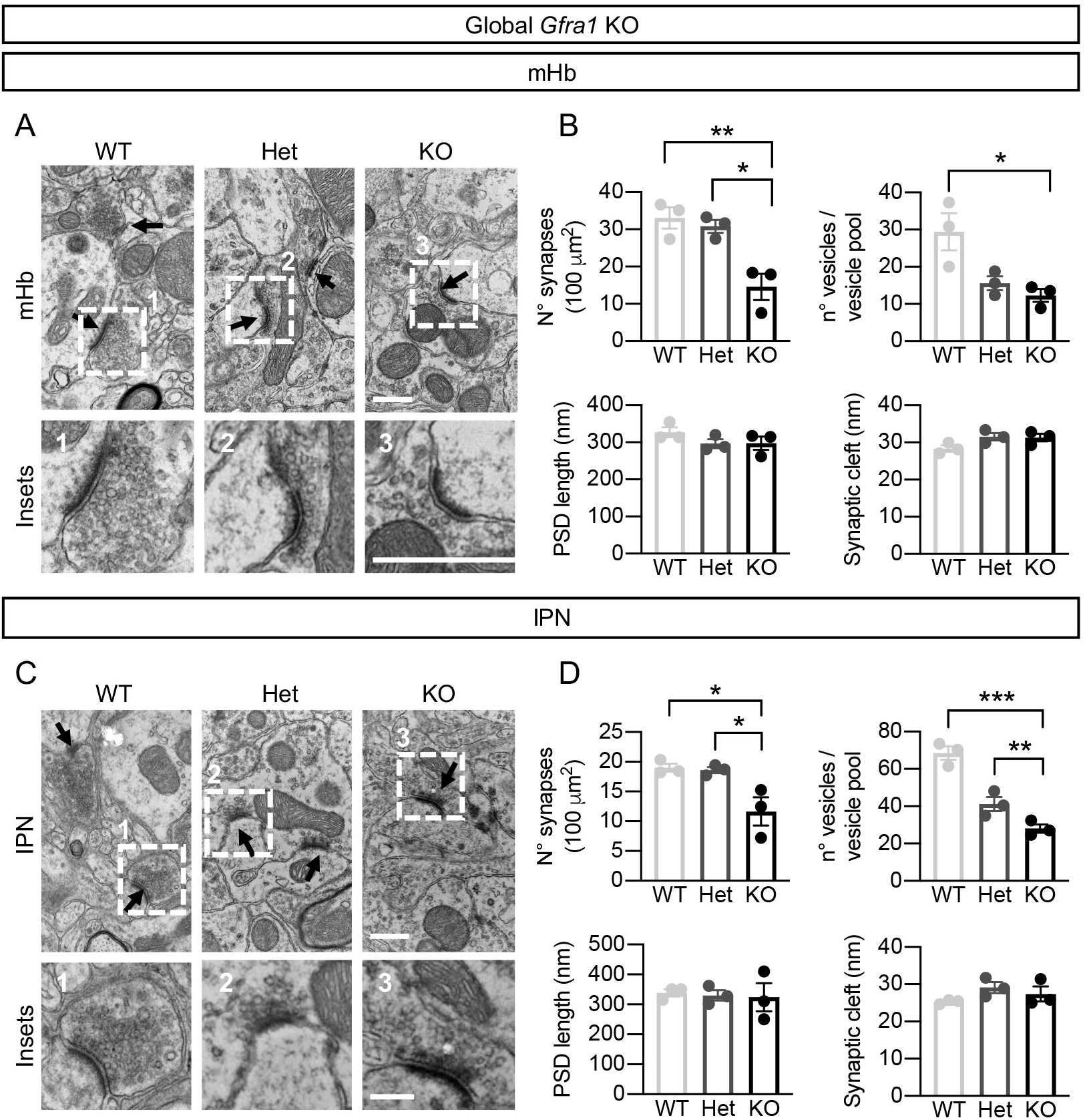
Reduced number of synapses and presynaptic vesicles in mHb and IPN after acute ablation of GFRα1 in adult mice. A, C) Transmission electron microscopy (TEM) images of the mHb (A) and the IPN (C) of WT, Het and KO mice. Scale bars, 500nm and 200nm (insets). B, D) Quantification (±SEM) of synapse density, synaptic cleft width, number of presynaptic vesicles and postsynaptic density (PSD) length in mHb (B) and IPN (D) of WT, Het and KO mice. N=3 mice per group; 1-way ANOVA followed by Tukey’s post-hoc test; *, P< 0.05; **, P< 0.01; ***, P< 0.001).

### Adult loss of GFRα1 affects the functional connectivity of the septum→Hb→IPN pathway and alters AMPAR-mediated glutamatergic responses in mHb neurons

In order to assess whether adult loss of GFRα1 affects connectivity in the septum→mHb→IPN pathway, we used a chemogenetic approach. In the first set of experiments (Fig. 4A), an AAV2 vector containing a DREADD (Designer Receptor Exclusively Activated by Designer Drug) construct driving Cre-dependent expression of hM3Dq and mCherry was stereotaxically injected in the TS of WT, Het and KO mice (Fig. 4A). Since the TS neurons projecting to the mHb do not express GFRα1 (i.e. no Cre expression in Het or KO), a second AAV2 construct expressing both Cre and GFP was delivered together with the DREADD construct. Additional control experiments showed that all genotypes were infected by the AAV2 vectors with comparable efficiency (Fig. S5A). Treatment with clozapine-N-oxide (CNO) for one hour resulted in strong c-Fos signals in the mHb only when the DREADD construct was expressed by the septal neurons (Figs. 4B and S5B). Importantly, the number of c-Fos^+^ cells was significantly reduced in both Het and KO compared to WT (Fig. 4C), indicating that GFRα1 is required to maintain functional connectivity in the septum-mHb pathway.

**Fig 4.**
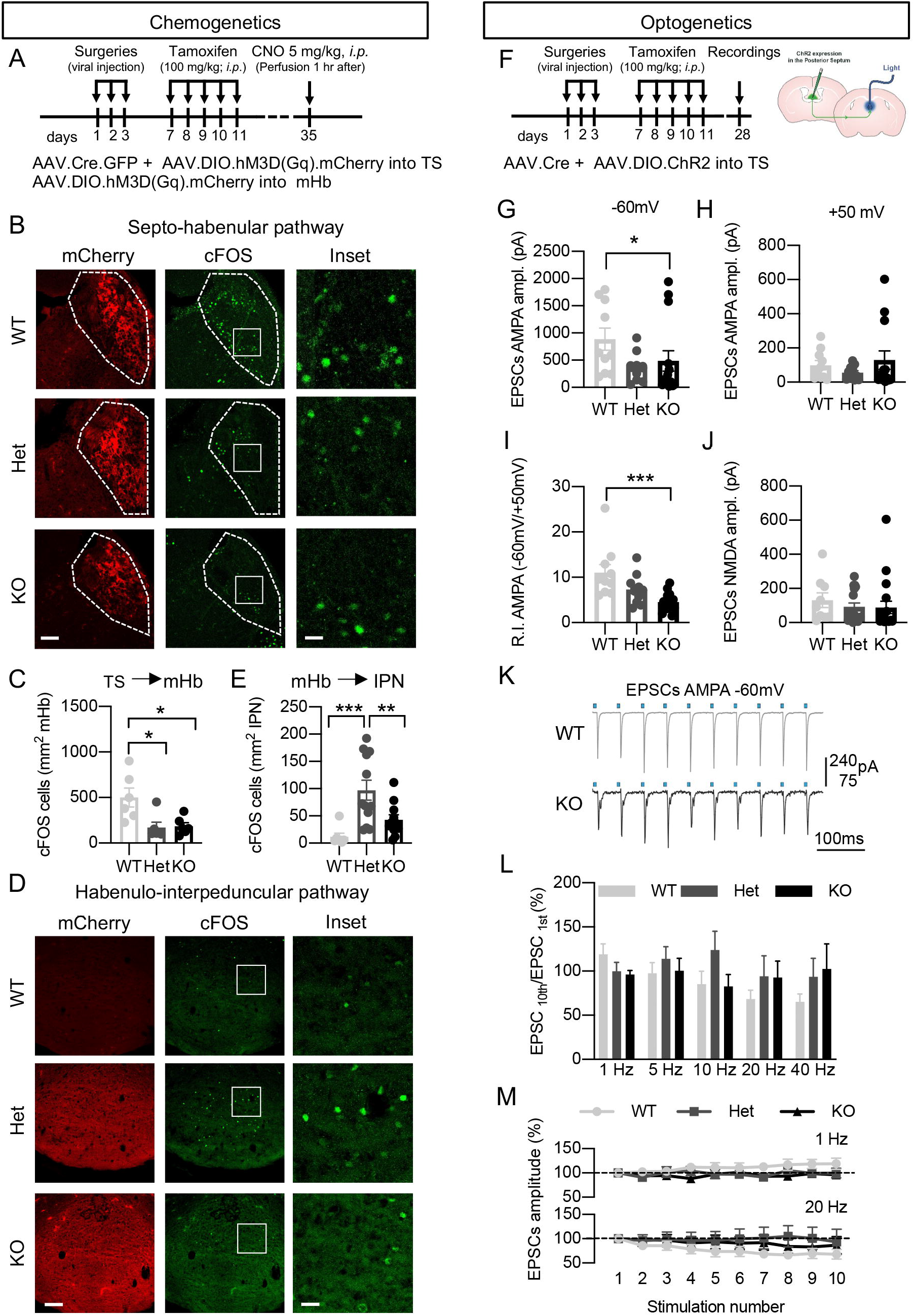
Adult loss of GFRα1 affects the functional connectivity of the septum→Hb→IPN pathway and alters AMPAR-mediated glutamatergic responses in mHb neurons. A) Schematics of the chemogenetic procedures. B, D) mCherry epifluorescence (red) and c-FOS (in green) immunostaining in the mHb (B) and IPN (D) of WT, Het and KO mice after chemogenetic activation of TS terminals (B) or mHb neurons (D) expressing hM3Dq. Scale bars, 70μm and 15 μm (insets) in (B), 100μm and 25 μm (insets) in (D).C, E) Quantification (±SEM) of c-FOS positive cells in mHb (C) and IPN (E) of WT, Het and KO mice. N=6 (C) and N=7-12 (E) mice per group (10-14 images (C) and 5-6 images (E) per mouse); 1-way ANOVA followed by Tukey’s multiple comparison test (C) or Bonferroni’s post hoc test (E); *, P< 0.05; **, P< 0.01; ***, P< 0.001. F) Schematics of the optogenetics procedures. G-I) EPSC average amplitudes at -60mV (G) and +50mV (H) mediated by AMPA receptors after optogenetic stimulation of septal fibers recorded in the presence of NMDA antagonist APV. Rectification index (I) was calculated as the ratio between the EPSC amplitudes at -60mV and +50 mV. N=10-14 cells per genotype; Kruskal-Wallis followed by Dunn’s multiple comparison test; *, P< 0.05; ***, P< 0.001. J) NMDAR-mediated EPSCs at +50mV in WT (n=9), Het (n=17) and KO (n=18) mHb cells after optogenetic stimulation of the septal terminals. K) AMPAR-mediated EPSCs recorded at -60mV in mHb of WT and KO mice during 10 consecutive optogenetic 20Hz stimulations. L, M) EPSC amplitude (±SEM) (L) and its variation (M) between 1^st^ and 10^th^ stimulation at different frequencies in mHb of WT, Het and KO mice.

In the second set of experiments, the same DREADD virus was injected in the mHb, while a second AAV2 virus expressing only GFP was used as control. In this case, only Het and KO mice express Cre (from the *Gfra1* locus), thus in WT mice no hM3Dq expression could be detected in the mHb (Fig. S5C) neither c-Fos signals in the IPN after CNO treatment. c-Fos induction could neither be detected in Het or KO mice when either tamoxifen or CNO were omitted (Figs. S5D, E). Treatment with CNO induced strong c-Fos signals in the IPN only in the presence of the DREADD virus (Figs. 4D and S5F). In this case, the number of c-Fos^+^ cells was significantly reduced in the IPN of KO mice compared to Het mice (Fig, 4E), indicating impaired connectivity in the adult mHb-IPN pathway after acute loss of GFRα1.

The c-Fos responses observed in the mHb reflected the result of sustained chemogenetic stimulation (one hour CNO treatment) of TS neurons. The decrease observed in Het and KO mice is in line with their reduced number of presynaptic vesicles compared to WT animals, as a smaller vesicle pool would be faster depleted following prolonged activation. In order to investigate faster synaptic responses, we performed electrophysiological measurements of AMPAR- and NMDAR-mediated components in the mHb after optogenetic stimulation of TS afferents. AAV2 vectors expressing Cre-dependent Channelrhodopsin-2 (ChR2) or Cre recombinase were stereotaxically co-injected in the TS of WT, Het and KO mice (Fig. 4F). Brain slices containing the mHb were prepared from 3 weeks after tamoxifen treatment, and excitatory post-synaptic currents (EPSCs) elicited by optogenetic stimulation of TS terminals were recorded in voltage-clamped mHb neurons. These EPSCs could be completely blocked by the AMPAR and NMDAR antagonists NBQX and D-APV, respectively, (data not shown), indicating that they were purely glutamatergic, in agreement with previous studies (11). We found that the amplitude of AMPAR-mediated EPSCs recorded at -60mV in the presence of the NMDAR antagonist D-APV was significantly reduced in mHb neurons of KO mice compared to WT (Fig. 4G). On the other hand, no difference between genotypes could be found in EPSCs recorded at +50mV (Fig. 4H). As a result, the ratio between EPSCs amplitudes measured at these two voltages, which constitutes the rectification index (R.I.) of the AMPAR component of these EPSCs, was significantly lower in mHb KO neurons compared to WT neurons (Fig. 4I). Neither AMPAR-mediated EPSCs amplitude nor R.I. of Het neurons were significantly different from WT (Figs. 4G-I). Importantly, NMDAR-mediated EPSCs were not altered in KO mHb neurons (Fig. 4J). Neither the dynamic behavior of the synapses in response to trains of 10 stimulations at frequencies ranging from 1 to 40 Hz was different between KO and WT mHb neurons (Figs. 4K-M), suggesting that the differences observed between genotypes in EPSCs amplitude were due to postsynaptic alterations. We note that the presence of rectifying AMPAR-mediated EPSCs (Fig. 4I) is evidence of the existence of Ca^2+^-permeable AMPARs in mHb neurons, in agreement with previous observations (Otsu et al., 2018). Thus, the lower R.I. value of AMPAR-mediated EPSCs in KO mHb neurons indicated reduced Ca^2+^ permeability of AMPAR complexes in mHb neurons upon removal of GFRα1, suggesting possible changes in the composition of postsynaptic AMPAR subunits in these neurons.

### Altered molecular composition of AMPAR complexes in mHb neurons after adult ablation of GFRα1

AMPAR are tetrameric ion channels formed by dimers of dimers derived from any of four different subunits, termed GluA1 to 4, in different combinations (31). In the adult brain, the majority of AMPAR complexes contain GluA2 subunits that have undergone RNA editing, making the assembled channel Ca^+2^-impermeable (32). On the other hand, AMPAR lacking GluA2 are Ca^+2^-permeable and, although very abundant in the developing CNS, they are not common in the adult (31). Although earlier electrophysiological data have indicated the presence of Ca^2+^-permeable AMPAR complexes in the septo-habenular-interpeduncular pathway (9,11), the molecular composition and functional importance of these complexes are unknown. Immunohistochemistry for each of the four GluA subunits in the mHb of adult WT mice revealed high levels of GluA1 and GluA4, lower expression of GluA2, and negligible expression of GluA3 (Fig. S6A). In the IPN, all GluA subunits were present, with surprisingly high levels of GluA4 (Fig. S6A). Western-blot analysis of whole tissue, cytosolic and synaptic fractions of the mHb and IPN revealed the presence of all four subunits in both structures (Fig. S6B). While the GluA3 blot had to be overexposed to detect specific signals, GluA4 was again very highly expressed in the IPN (Fig. S6B). To better understand the composition of AMPAR complexes in the mHb and IPN, the Proximity Ligation Assay (PLA) was used to investigate interactions *in situ* between GluA1 and GluA2 subunits, the most common Ca^+2^-impermeable channels in the brain, and between GluA1 and GluA4, forming Ca^+2^-permeable complexes. Both types of complexes were equally present in the dorsal and the ventral portions of the mHb (Figs. S6C, D), while in the IPN the lateral subnuclei presented less GluA1/GluA2 and GluA1/GluA4 complexes than the core IPN (Figs. S6C, E).

Having established the presence of both GluA1/GluA2 (Ca^2+^-impermeable) as well as GluA1/GluA4 (Ca^2+^-permeable) AMPAR complexes in mHb and IPN, we examined possible changes in the levels, arrangement and phosphorylation state of these subunits upon removal of GFRα1, as any of these can lead to changes in synaptic strength and function (33). In the mHb, the level of GluA2 was significantly increased in KO mice (Fig. 5A), while in the IPN, both GluA1 and GluA4 levels were reduced in mice lacking GFRα1 (Fig. 5B). No statistically significant changes were observed in Het mice. Whereas GluA1 phosphorylation at Ser^831^ was unchanged in the mutants, phosphorylation at Ser^845^ was reduced in the mHb of KO mice (Figs. 5C-F). Phosphorylation at Ser^845^ has been implicated in the insertion of GluA1 into synapses (34). Changes in the composition of AMPAR complexes were evaluated by PLA. GluA1/GluA2 interactions were found to be significantly increased in the mHb and core IPN of KO mice (Figs. 5G, H), while GluA1/GluA4 complexes were decreased in the core IPN of the mutants (Figs. 5I, J), indicating that adult loss of GFRα1 leads to rearrangements in the composition of postsynaptic AMPAR subunits in the septo-habenular-interpeduncular pathway which are consistent with the reduced Ca^2+^ permeability of those channels.

**Fig 5.**
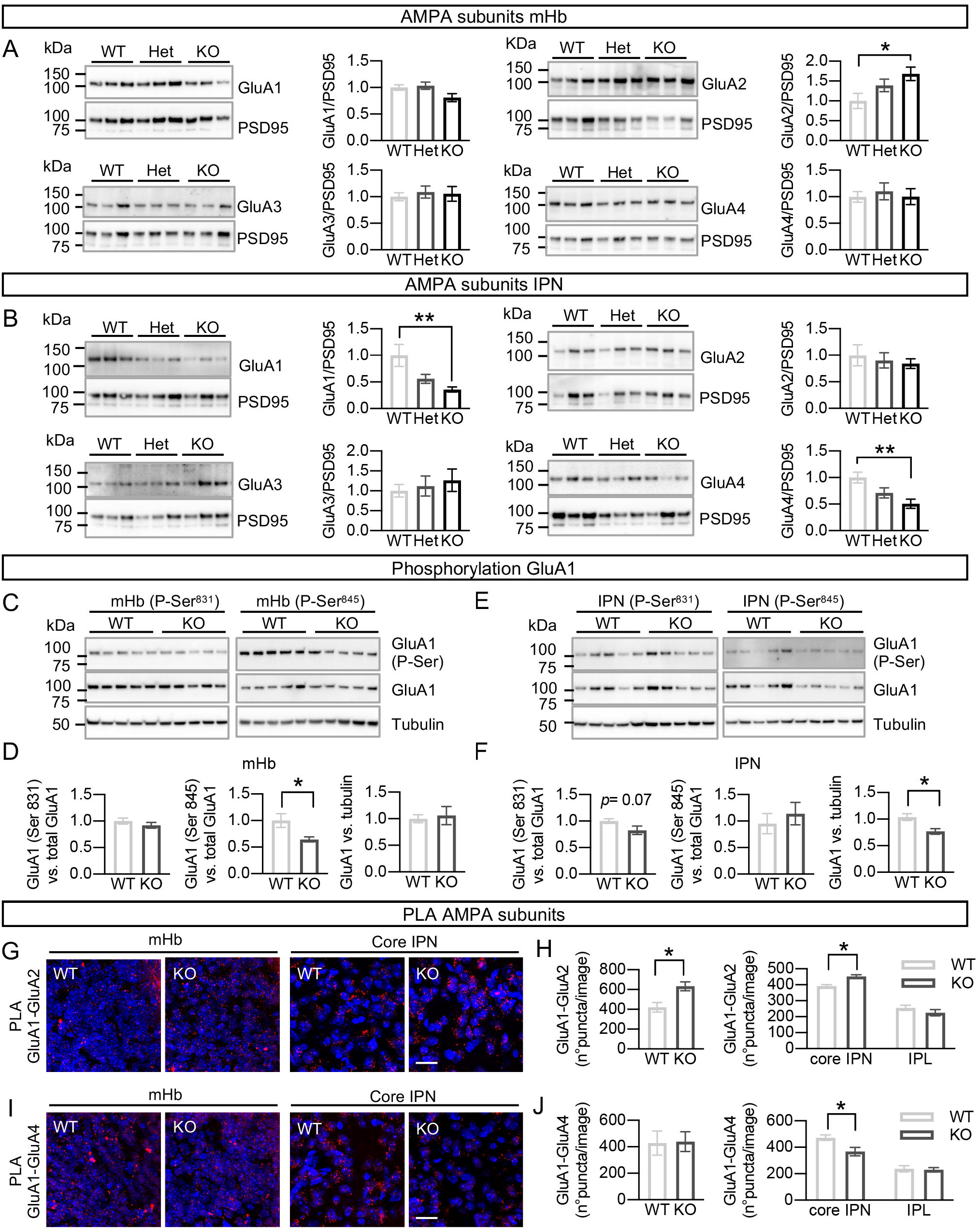
Altered molecular composition of AMPAR complexes in mHb neurons after adult ablation of GFRα1. A, B) Immunoblots of synaptic protein fractions from mHb (A) and IPN (B) of WT, Het and KO mice probed for GluA1, 2, 3 and 4 as indicated. PSD95 was probed as loading control. Quantifications (±SEM) of GluA1 to 4 levels were corrected for PSD95 levels and normalized to levels in WT samples. N=10 mice per group; 1-way ANOVA analysis followed by Tukey’s multiple comparison test; *, P<0.05; **, P< 0.01. C, E) Immunoblots of synaptic protein fractions from mHb (C) and IPN (E) of WT, Het and KO mice probed for GluA1 P-Ser^831^ and P-Ser^845^ as indicated, and reported for total GluA1 and Tubulin as loading controls. D, F) Quantifications (±SEM) of GluA1 P-Ser^831^ and P-Ser^845^ levels in mHb (D) and IPN (F) corrected for total GluA1 levels and normalized to levels in WT samples. N=13 (D) and 14-15 (F) mice per group; Student’s T-test; *, P<0.05 G, I) Proximity Ligation Assay (PLA) signals (red) for GluA1-GluA2 (G) and GluA1-GluA4 (I) complexes in coronal sections of mHb and IPN from WT and KO mice. Counter-staining with DAPI appears in blue. Scale bars, 20 μm. H, J) Quantification (±SEM) of PLA puncta for GluA1-GluA2 (H) and GluA1-GluA4 (J) complexes in mHb, core IPN and lateral IPN (IPL) from WT and KO mice. N=5-6 mice per group (total 24 images per group); Student’s T-test; *, P<0.05.

### Acute loss of GFRα1 in adult mHb neurons reduces anxiety-like behavior and potentiates context-based fear responses

Selective lesion of the TS and BAC projections to the mHb has been shown to increase fear responses to electric shock while decreasing anxiety-like behavior (4). We performed a battery of behavioral tests 2 weeks after tamoxifen administration in WT, Het and KO mice or 2 weeks after Cre virus injection in mHb.KO mice. The open field and elevated plus maze were used to evaluate anxiety; foot shock-induced freezing behavior, to evaluate innate fear responses; and passive avoidance and fear conditioning, to evaluate fear-based learning. General locomotor activity in the open field was unchanged among genotypes, although KO mice showed a trend towards increased distance and time spent in the center of the arena, which did not reach statistical significance (Fig. S7A). In the elevated plus maze, KO mice entered more times and spent significantly longer time in the open arms than WT and Het animals, suggesting reduced anxiety-like behavior in adult mice lacking GFRα1 (Fig. 6A). Importantly, similar results were obtained in mHb.KO mice after Cre virus injection to the mHb (Fig. S7B and Fig. 6B), indicating that GFRα1 is required in the adult mHb for normal anxiety-like behaviour.

**Fig 6.**
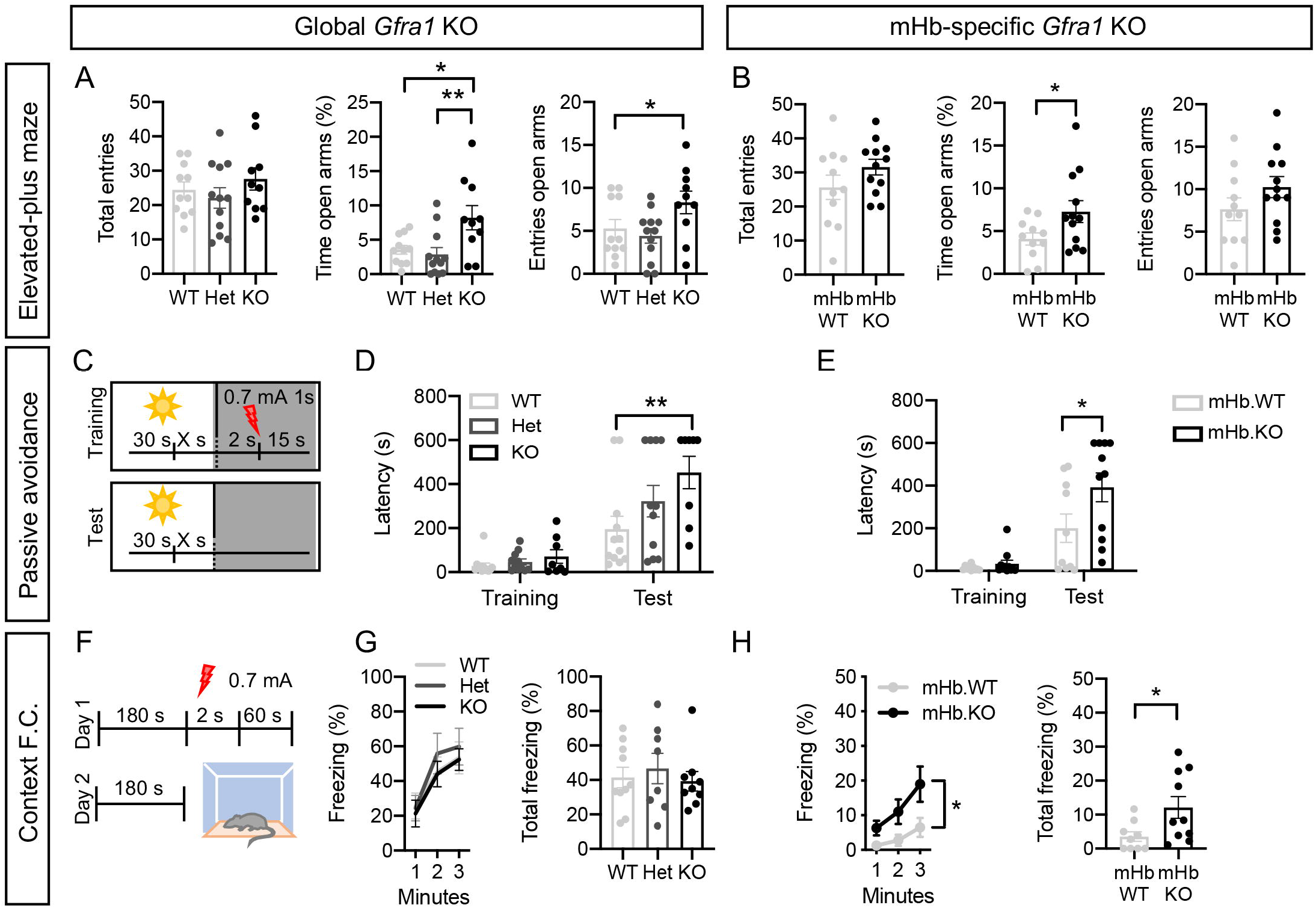
Acute loss of GFRα1 in adult mHb neurons reduces anxiety-like behavior and potentiates context-based fear responses. A, B) Total entries, time spent in open arms and open arms entries (±SEM) for WT, Het and KO mice (A) and mHb.WT and mHb.KO mice (B) in the Elevated Plus Maze test. N=10-12 mice per group; 1-way ANOVA followed by Bonferroni post-hoc test (A) and Student’s t-test (B); *, P<0.05; **, P<0.01. C-E) Schematic (C) and latencies (±SEM) during training and test for WT, Het and KO mice (D) and mHb.WT and mHb.KO mice (E) in the Passive Avoidance test. N=8-12 mice per group; 2-way ANOVA followed by Bonferroni post-hoc test; *, P<0.05; **, P<0.01. F-H) Schematic (F), freezing time-course and total freezing time (±SEM) for WT, Het and KO mice (G) and mHb.WT and mHb.KO mice (H) in the Context Fear Conditioning test. N=8-10 mice per group; 2-way ANOVA followed by Bonferroni post-hoc test and Student’s t-test; *, P<0.05; **, P<0.01.

Freezing responses after three consecutive foot shocks were similar between WT, Het and KO mice (Fig. S7C) as well as mHb.KO and mHb.WT animals (Fig. S7D), indicating unaltered innate fear behavior in this test after loss of GFRα1. In the passive avoidance test (Fig. 6C), both global (KO) and mHb-specific (mHb.KO) ablation of GFRα1 significantly increased the avoidance latency (Figs. 6D, E), indicating potentiated responses to fear-based learning after acute loss of habenular GFRα1. To assess fear conditioning, mice were trained to pair a foot-shock with either a specific context (Fig. 6F) or a tone cue (Fig. S7E). In the context fear conditioning, there were no differences between the WT, Het and KO groups (Fig. 6G), but mHb.KO mice lacking GFRα1 only in mHb neurons showed an exacerbated freezing response after conditioning compared to mHb.WT controls (Fig. 6H). Finally, cued conditioned fear responses did not differ between WT, Het and KO mice, or between mHb.KO and mHb.WT mice (Figs. S7F, G). Together, the results of these studies show that context-conditioned fear responses are significantly enhanced after acute loss of GFRα1 in adult mHb neurons.

## Discussion

Fear is an evolutionary conserved, adaptive behavior that helps organisms to react to threatening or otherwise dangerous situations. On the other hand, persistent or exaggerated fear in the absence of danger is detrimental and maladaptive. Previous research has shown that neurons in the mHb play a critical role in calibrating behavioral responses to threatening situations in an experience-dependent manner (4,5). mHb neurons display a number of features that set them apart from the majority of neurons in the adult brain, including excitatory GABAergic inputs and Ca^2+^-permeable AMPA receptors. However, the mechanisms that maintain and regulate the unique properties of mHb neurons and their function in mediating fear responses remain incompletely understood. Here, we demonstrate that the neurotrophic receptor GFRα1 regulates synapse homeostasis in adult mHb neurons to facilitate the normal processing of fear behavior. We report three major findings. First, GFRα1 is highly abundant in adult mHb neurons and their terminals, where it is localized to synapses, and required for the maintenance, structural integrity and function of septo-habenular and habenulo-interpeduncular synaptic connections. Second, GFRα1 favors the expression of Ca^2+^-permeable AMPA receptor complexes formed by the association of GluA1 and GluA4 subunits, at the expense of Ca^2+^-impermeable GluA1/GluA2 complexes, in adult mHb synapses. Third, acute loss of GFRα1 in adult mHb neurons reduces anxiety-like behavior and potentiates context-based fear responses, phenocopying the effects of lesions to mHb afferents from the posterior septum. These findings establish an unexpected role for GFRα1 as a critical mediator of the stability and function of glutamatergic synapses in the adult brain.

GDNF was originally discovered as a survival factor for midbrain dopaminergic neurons and a candidate anti-Parkinson’s Disease agent (35), thereafter focusing research on GDNF receptors on the SNpc and VTA areas that harbor these cells. More recent studies characterized other sites of GFRα1 expression in the mammalian brain, particularly in GABAergic neurons of the cerebral cortex (36,37), cerebellum (38,39) and olfactory bulb (40,41). Our present findings establish mHb neurons as the cells that express the highest levels of GFRα1 in the adult mouse brain, higher than any other known site of expression, and the first glutamatergic neurons known to require this receptor for appropriate connectivity and function. Although extensively studied in the developing nervous system, the functions of neurotrophic systems in the adult brain are less well understood. With the possible exception of neural injury and neurodegenerative disease, adult deletion of neurotrophic factors or their receptors has in most cases no effect on neuron maintenance. Our discovery of the requirement of GFRα1 for the structural and functional stability of mHb glutamatergic synapses represents the first demonstration of a function for this receptor in mature adult neurons. Despite being only expressed by mHb neurons, GFRα1 was found enriched at both septo-habenular and habenulo-interpeduncular synapses, suggesting both pre and post-synaptic functions. This may be possible through the release of GFRα1 from the cell surface (13,14), facilitating non-cell-autonomous effects across the synapse and peri-synaptic areas. In contrast to RET, expression of NCAM was detected in neurons along the septum→Hb→IPN pathway, where it was also concentrated at synapses, making it a possible candidate for mediating the effects of GFRα1 in this circuit. NCAM has been found to contribute to several functions in synapse development and maintenance beyond cell adhesion, including spine formation (42), synapse maturation (43), presynaptic vesicle recycling (44), and NMDA- and AMPA-mediated glutamatergic signaling (45). GDNF, the main ligand of GFRα1, could also be detected in the septum, Hb and IPN, although its specific cellular source, e.g. neurons or glia, remains to be established. The specific contributions, if any, of NCAM and GDNF to the function of GFRα1 in the septum→Hb→IPN pathway will require further investigation. On the other hand, the abundance of GFRα1 protein in cell bodies and terminals of mHb neurons suggests other mechanisms may also be possible, including structural roles in overall synapse stability, and regulation of trafficking or turn-over of AMPAR subunits.

Our behavioral studies in adult mice following acute loss of GFRα1 in mHb neurons revealed an unexpected requirement for this receptor in normal anxiety-like behavior and conditioned fear responses. The behavioral effects observed in the mutants resembled lesions to glutamatergic TS and BAC afferents to the mHb (4), indicating that GFRα1 is a positive regulator of glutamatergic transmission in the septum→Hb→IPN pathway, and an essential component for normal mHb neuron function. The haploinsufficiency observed in HET mice for some of the phenotypes indicates a dosage effect, and suggests a potential mechanism for adjusting the functional response of the pathway to dangerous or threatening situations by regulating the levels of GFRα1, its ligands or co-receptors. In this context, it would be important to identify the environmental cues and/or internal states that may affect GFRα1 expression in mHb neurons, as well as its trafficking and accumulation in mHb terminals. Finally, our findings open up a new avenue for investigations into the functional plasticity of mHb neurons, which will not only improve our understanding of psychiatric disorders associated with fear and anxiety, but also potentially lead to novel treatments for such disorders.

## Methods

### Mice

Mice were housed 2-5 per cage in standard conditions in a temperature and humidity-controlled environment, on a 12/12 h light/dark cycle with *ad libitum* access to food and water. The following transgenic mouse lines were used for experiments: *EIIa^Cre^* (46), *Gad67^Cre^* (27), *Gfra1^Cre-ERT2^* (38), *Gfra1^LoxP^*(kindly provided by M. Saarma and J.O Andressoo, University of Helsinki), *R1CG^LoxP^,* which expresses GFP from the *gfra1* locus after Cre-mediated recombination (26), *Ret^GFP^* (24) *Rosa26^dTom^* (47). The *gfra1^gfp^* mouse line was generated by crossing *EIIa^Cre^* with *R1CG^LoxP^* mice to obtain GFP expression under the *Gfra1* promoter in the germ cells and subsequently crossing the offspring with C57BL/6 mice. All animals were bred in a C57BL6/J background (Jackson) with the exception of the *gfra1^LoxP^* that was kept in a CD1 background (Jackson). Therefore, all animals resulting from breeding *Gfra1^Cre-ERT2^* and *Gfra1^LoxP^* mice were 50:50 C57BL6/J: CD1 background. Both males and females were used for these studies. Animal protocols were approved by Stockholms Norra Djurförsöksetiska Nämnd and are in accordance with the ethical guidelines of Karolinska Institutet (permit numbers N173-15; 11563-2018 and 11571-2018).

### Tamoxifen treatment and stereotaxic surgery

Adult mice (3-4 months old) were injected intraperitoneally (i.p.) with 100 mg/kg of tamoxifen (Sigma-Aldrich) dissolved in 10% EtOH in corn oil (Sigma-Aldrich) for 5 consecutive days. Adult mice (3-4 months old) were anesthetized with isoflurane (1- 5%) and placed in a stereotaxic frame (David Kopf Instruments). Injection was performed with a wiretrol capillary micropipette (Drummond scientific) by nanoliter pressure injection at a flow rate of 50 nL per min (Micro4 controller, World Precision Instruments; Nanojector II, Drummond Scientific). The pipette was left on place for 10 min after the injection before retracting it slowly from the brain. *Gfra1^LoxP^* mice were injected bilaterally with 300 nL of AAV2-CMV-PI-Cre-rBG (Penn Vector Core, 2.27×10^13^ genomic copies per mL) or vehicle in the following coordinates obtained from the Paxinos and Franklin atlas: (i) mHb (10° angle at the coordinates anteroposterior AP -1.94 mm, mediolateral ML ±0.75 mm, dorsoventral DV –2.3 mm). Due to the lack of a reporter gene in the viral construct, the expression levels of GFRa1 in the mHb were evaluated in all cases by immunofluorescence at the end of the experiment and the ones showing no loss were removed from the experimental data set. To evaluate the transfection efficiency of the viral vector, *R1CG^LoxP^* mice were injected with the virus in the same coordinates, observing GFP expression under the *gfra1* promoter after Cre recombination.

### Biochemical techniques

#### Tissue processing

Animals were sacrificed by cervical dislocation, brains were rapidly removed, dissected manually, frozen in dry ice and stored at -80 °C until further use. For the dissection of the mHb and the IPN, brains were cut coronally with a blade at -1mm from bregma and +1mm from lambda, obtaining a coronal section containing the habenula. The cortex and the hippocampus from that section were removed, leaving the mHb on sight, which was dissected by pulling it out carefully. Next the IPN, which could be visualized directly in the remaining caudal part of the brain, was collected.

#### RNA extraction and Real-time RT-PCR

The mRNA levels of *Gfra1* and *Gdnf* were quantified by real-time RT-PCR in total RNA collected from the brain areas of interest relative to a geometric mean of mRNAs encoding the ribosomal protein 18S. Total mRNA was isolated from the dissected brain areas using the RNeasy Mini Kit (Qiagen) according to manufacturer’s protocol. The purity and quantity of RNA were measured in a nanodrop 1000 (ThermoScientific). cDNA was synthesized by reverse transcription using 150 ng of RNA in a reaction volume of 20 ul using the High Capacity cDNA reverse transcription kit (Applied Biosystems) according to the manufacturer’s protocol. cDNA samples were diluted 1:3 in H2O before used, and to measure the levels of 18S a further 2:800 dilution was made. Real-time RT-PCR was conducted using the 7500 Real-Time PCR system (Applied Biosystems) with SYBR Green fluorescent probes using the following conditions: 40 cycles of 95°C for 15 s, 60°C for 1 min and 72°C for 30s. Forward and reverse primers were ordered from Sigma and used at a concentration of 100 nM each. The following primer pairs were use: gfra1 (Fw: 5’- GAAGATTGCCTGAAGTTTCTGAAT-3’; Rv: 5’-GGTCACATCCGAGCCATT-3’); gdnf (Fw: 5’-GTGACTCCAATATGCCTGAAGA-3’; Rv: 5’-CGCTTGTTTATCTGGTGACCT- 3’); 18S (Fw: 5’- CAATTATTCCCCATGAACG-3’; Rv: 5’- GGCCTCACTAAACCATCCAA-3’). Five biological replicates were performed for each study group. All the experiments were carried out in triplicates for each data point. Data analysis was performed using the 2−ΔCt method, and relative expression levels were calculated for each sample after normalization against the housekeeping gene 18S.

#### Protein collection and Western-blot

The whole extract and the cytosolic and synaptosome protein fractions were collected using the Syn-PER™ Synaptic Protein Extraction Reagent (ThermoFisher) following manufacture instructions. The mHb and the IPN of two mice per genotype were pulled together due to the small size of the nuclei of interest. Protein concentration was measured with the Pierce™ BCA Protein Assay Kit (ThermoFisher) and samples were prepared for SDS-PAGE (20 ug per sample) in SDS sample buffer (Life Technologies) and boiled at 95°C for 10 min before electrophoresis on 4-12% pre-cast gels (Sigma) following manufacture instructions. Proteins were transferred to polyvinylidene fluoride (PVDF) membranes (Amersham). Membranes were blocked with 5% non-fat milk in TBST and incubated with primary antibodies overnight in 1% milk. The following primary antibodies were used at the indicated dilutions: Goat anti GFRα1 (1:500, AF560, RnD): goat anti c-RET (1:500, AF482, RnD); rabbit anti NCAM (1:1000, AB5032, Millipore); Mouse anti PSD95 (1:1000, MA1-046, Thermo Scientific); Mouse anti GDNF (1:200; Sc-13147, Santa Cruz); rabbit anti GluA1 (1:1000, AB1504, Millipore); rabbit anti GluA2 (1:1000, AB1768-I, Millipore); rabbit anti GluA3 (1:1000, AGC-010, Alomone Labs); rabbit anti GluA4 (1:1000, AB1508, Millipore); rabbit anti phospho-Ser831 GluA1 (1:500, 75574, Cell Signaling); rabbit anti phospho-Ser845 GluA1 (1:500, 8084, Cell Signaling); mouse anti tubulin alpha (1:10000, T6199, Sigma). Immunoreactivity was visualized using appropriate horseradish peroxidase (HRP)-conjugated secondary antibodies (1:5000, Dako). Immunoblots were developed using the ECL Advance Western Blotting Detection kit (Life Technologies) or the SuperSignal West Femto Maximum sensitivity substrate (ThermoFisher) and images were acquired with the Image-Quant LAS4000 (GE Healthcare). Image analysis and quantification of band intensities were done with ImageQuant software (GE Healthcare).

### Histological techniques

#### Tissue processing

Mice were deeply anaesthetized with isoflurane (Baxter Medical AB), and transcardially perfused with 25 ml of 0.125 M phosphate buffered saline (PBS, pH 7.4, Gibco) and 40 ml of 4% paraformaldehyde (PFA, Histolab Products AB). Brains were removed, post fixed in PFA overnight and cryoprotected in 30% sucrose in PBS. To obtain 20 μm floating sections, brains were embedded in OCT compound (Sakura), frozen at -80°C and cut with a cryostat (CryStarNX70, ThermoScientific). To obtain 30 μm sections, brains were cut in a microtome (Leica SM2000 R). All sections were stored at -20°C in a cryoprotective solution containing 1% DMSO (Sigma-Aldrich) and 20% glycerol (Sigma-Aldrich) in 0.05M Tris-HCl pH 7.4 (Sigma-Aldrich).

#### General immunofluorescence

30 μm brain sections were washed in PBS and blocked for 1 h in 5% normal donkey serum (NDS, Jackson Immunoresearch) and 0.3% Triton X-100 (Sigma-Aldrich) in PBS. Incubation with the primary antibodies was done overnight at 4°C in blocking solution. After 3×10 min washes with PBS, the sections were incubated with the corresponding secondary donkey Alexa fluorescent antibodies (1:1000, ThermoFisher) and 0.1 mg/ml of 40-6-diamidino-2-phenylindole (DAPI; Sigma-Aldrich) for 2h at RT. Sections were finally washed with PBS 3×10 minutes, mounted on glass slides in a 0.2% solution of gelatin (Sigma-Aldrich) in 0.05M Tris-HCl buffer (pH 7.4), dried and cover slipped in DAKO fluorescent mounting medium. The primary antibodies used in this study were: Goat anti GFRα1 (1:500, AF560, RnD); Goat anti ChAT (1:500, AP144P, Millipore); Chicken anti GFP (1:500, ab13970, Abcam); rabbit anti NCAM (1:1000, AB5032, Millipore); rabbit anti cFOS (1:500, sc-52, Santa Cruz); rabbit anti calretinin (1:500, AB5054, Chemicon); mouse anti calretinin (1:500, 6B3, SWANT); rabbit anti GluA1 (1:500, AB1504, Millipore); rabbit anti GluA2 (1:500, AB1768-I, Millipore); rabbit anti GluA3 (1:500, AGC-010, Alomone Labs); rabbit anti GluA4 (1:500, AB1508, Millipore); rat anti substance P (1:500, Millipore, MAB356).

#### Retrograde tracing

300 nL of fluorogold (Fluorochrome) were injected unilaterally in the mHb of *Gfra1^GFP^* and *Ret^GFP^* adult mice at the following coordinates obtained from the Paxinos and Franklin atlas: 10° angle, AP -1.94 mm, ML +0.75 mm, dorsoventral DV –2.3 mm. Mice were perfused 2 weeks later.

#### Immunofluorescence for synaptic markers

20 μm brain sections were washed in PBS, blocked for 1hr at RT in 20% NDS in PBS, incubated with the primary antibodies at 4°C overnight in a solution containing 10% NDS and 0.3% TX-100, washed 3×1hr in PBS, incubated in the same solution as the primary antibodies with the corresponding secondary donkey Alexa fluorescent antibodies (1:1000, ThermoFisher) for 2hr, washed 3×1hr in PBS, counterstained with DAPI for 10 min and washed for 2×30 min in PBS. Sections were finally mounted on glass slides in a 0.2% solution of gelatin (Sigma-Aldrich) in 0.05M Tris-HCl buffer (pH 7.4), dried and cover slipped in DAKO fluorescent mounting medium. The primary antibodies used were guinea pig anti VGlut1 (1:500, AB2251, Millipore); Mouse anti VGlut2 (1:500, 135421, SYSY); rabbit anti PSD93 (1:500, 124102, SYSY); Goat anti GFRα1 (1:500, AF560, RnD); guinea pig anti VGAT (1:500, 131004, SYSY); Mouse anti gephyrin (1:500, 147111, SYSY); goat anti synapsin Ia/b (1:500, sc8295, Santa Cruz).

#### Proximity ligation assay (PLA)

PLA was performed according to manufacturer’s instructions (Duolink; Sigma-Aldrich). Briefly, 30 μm coronal sections containing the mHb and the IPN were blocked and permeabilized in 5% NDS and 0.3% TX-100 in PBS for 1h and incubated with rabbit anti GluA1 (1:400, AB1504, Millipore) and mouse anti GluA2 (1:400, Abcam, ab192760) or rabbit anti GluA1 (1:400, AB1504, Millipore) and goat anti GluA4 (1:400, ab115322, Abcam) overnight at 4°C. Mouse anti NMDAR2B (1:400, 610416, BD) and mouse anti DARP32 (1:400, sc-271111, Santa Cruz) were used as negative controls. Then sections were washed with PBS, incubated with minus and plus PLA probes for 1 hr at 37°C, washed with buffer A, incubated with Ligation-Ligase solution for 30 min at 37°C, washed again with buffer A, and finally incubated with the Amplification-Polymerase solution (Duolink™ In Situ detection reagents Orange) for 100 min at 37°C. After washing with buffer B, sections were incubated for 10 min with DAPI (1:10000 in buffer B), mounted on glass slides and coverslipped with DAKO mounting media.

#### Image acquisition and analysis

All fluorescent images were captured with a Carl Zeiss LSM 710 confocal microscope using ZEN2009 software (Carl Zeiss). All images from the same brain area and experiment were acquired using the same parameters. For imaging of the mHb and the IPN, one section every 120um was used for all experiments (6-8 sections per mHb and 4-5 sections per IPN). For the synaptic markers and the PLA experiments, 4 mHb images were obtained with the 63X objective per section (one dorsal and one ventral per mHb). For the IPN, 6 images were obtained per section (two dorsal, two ventral and two lateral). Image analysis was made with ImageJ and Fiji softwares. Quantification of the number of synaptic puncta and PLA dots was done in imageJ with the plugin synaptic puncta analyzer (48). In brief, background was subtracted from the image and the same intensity threshold and range of particle size was used to analyze all images from the same experiment.

#### Transmission electron microscopy

*Gfra1^+/+^*, *Gfra1^CreERT2/+^* and *Gfra1^Cre/fx^* 3-months old mice were treated with tamoxifen and perfused one month later with 2.5% glutaraldehyde and 1% Paraformaldehyde in 0.1M PB. Brains were removed and left on the same fixative overnight at 4°C. The day after 300 µm slices were obtained in a vibratome (Leica VT1000S) and the area corresponding to the mHb and the IPN were cut out. Subsequent tissue preparation and image acquisition was done at the electron microscopy unit (EMil) of Karolinska Institutet. The tissue blocks were washed in 0.1M PB; postfixed in 2% OsO4 in 0.1M PB for 2h at 4°C and then incubated with: (i) 70% EtOH for 30 min at 4°C; (ii) 95% EtOH for 30 min at 4°C; (iii) 100% EtOH with 0.5% uranil acetate for 20 min at RT; (iv) acetone 2×15 min RT; (v) LX-112/acetone (1:2) for 4h at RT; (vi) LX-112/acetone (1:1) overnight at RT; (vii) LX-112/acetone (1:2) for 8h at RT; (viii) LX-112 overnight at RT and finally embedded in LX-112 resin at 60°C. Ultrathin sections were cut with a EM UC6 (Leica) and 20 images/structure (dmHb, vmHb, IPR, IPV) from each mouse were acquired using both the FEI Tecnai 12 Spirit BioTwin transmission electron microscope (FEI Company, Eindhoven, The Netherlands) and the Hitachi HT7700 (Hitachi High-technologies) transmission electron microscope both operated at 100 kV and equipped with 2kx2k Veleta CCD cameras (Olympus Soft Imaging System).

### Chemogenetics and Optogenetics

#### Chemogenetics

*Gfra1^+/+^*, *gfra1^Cre/+^* and *gfra1^Cre/fx^* adult mice were stereotaxically injected: (i) In the posterior septum with 500 nL of a 1:1 mixture of AAV2-CMV-Cre-GFP (3.7×10^12^ genomic copies per mL; UNC Gene therapy vector core) and AAV2-hSyn-DIO- hM3D(Gq)-mCherry (6.1×10^12^ genomic copies per mL; UNC Gene therapy vector core) or the AAV2-CMV-Cre-GFP virus alone in the following coordinates obtained from the Paxinos and Franklin atlas :10° angle; AP -0.1 mm, ML +0.5 mm, DV – 2.58 mm. (ii) Bilaterally in the mHb with 300 nL of AAV2-hSyn-DIO-hM3D(Gq)-mCherry or 300nL of a control AAV2-CMV-GFP virus (5.4×10^12^ genomic copies per mL; UNC Gene therapy vector core) at the following coordinates: 10° angle, AP -1.94 mm, ML ±0.75 mm, DV –2.3 mm). One week after viral delivery animals were treated with tamoxifen to induce global ablation of *Gfra1* expression. Activation of hM3Dq was induced by an *i.p*. injection of clozapine-N-oxide (CNO, Tocris) dissolve in 2% DMSO in NaCl 0.9% serum (Braun) 28 days after the first dose of tamoxifen, and the animals were perfused 1h later.

#### Optogenetics

Ex vivo electrophysiological experiments were performed as described previously (11). Briefly, adult *gfra1^+/+^* and *gfra1^Cre/fx^* mice (3-4 months old) were injected with 200/300nL of a 1:1 mix of AAV2/1.EF1a.DIO.hChR2.eYFP and AAV2/9.CMV.Cre (with final titers of 2.02e13 and 4.90e12 GC/mL, respectively; Vector Core of the University of Pennsylvania, USA) in the posterior septum at the following coordinates, from bregma: ML 0.0 mm; AP 0.13–0.2 mm; DV -2.6 mm. Mice were treated with tamoxifen (4 consecutive daily *i.p.* injections, 100mg/kg) starting one week after surgery. Minimum 4 weeks following the injections, slices were obtained and whole-cell, voltage-clamp recordings of mHb neurons were performed at 32°C - 34°C in a bicarbonate-buffered saline containing (in mM): 116 NaCl, 2.5 KCl, 1.25 NaH_2_PO_4_, 26 NaHCO_3_, 30 glucose, 1.6 CaCl_2_, 1.5 MgCl_2_, and 5 × 10^−5^ minocycline (bubbled with 95% O_2_, 5% CO_2_), and using an intracellular solution containing (in mM): 135 CsMeSO_3_, 10 TEA-Cl, 4.6 MgCl_2_, 10 HEPES, 10 K_2_-creatine phosphate, 0.5 EGTA, 4 Na_2_-ATP, 0.4 Na_2_-GTP, 0.1 spermine and 1 QX-314, pH 7.35 and mOsm ∼300. Excitatory postsynaptic currents (EPSCs) were elicited by brief (2 to 4 ms-long) pulses of blue light provided by a 470-nm wavelength diode (Thorlabs, France) coupled to the slice chamber via the epifluorescence pathway of the microscope. EPSCs were recorded in the constant presence in the bath of the GABAa and glycine receptor blockers SR-95501 and strychnine respectively. Drugs were bath-applied: AMPAR antagonist NBQX (10µM), NMDA antagonist D-APV (50µM), SR-95501 (5µM) from Tocris Bioscience, and strychnine (2µM) from Sigma-Aldrich. Acquired with an EPC-10 double amplifier using the PatchMaster software (both from Heka Elektronik, Germany), the recordings were sampled at 10/40KHz, filtered at 10KHz and analyzed offline with custom routines written in Igor (Wavemetrics, USA). NMDAR-mediated EPSCs were quantified at +50mV, as the response amplitudes at the 7 ms-post peak time point, at full decay of the AMPAR component. The amplitudes of the AMPAR-mediated EPSC components at the holding potentials of +50mV and -60mV were measured at the peaks of the responses following bath application of the NMDA antagonist APV. Their ratio is the rectification index (R.I.) of the AMPAR currents. before D-APV application.

### Behavioral studies

All behavioral experiments were performed between 9 am and 2 pm, under low light conditions and blind to the genotype and treatment. The only sex differences found were on body weight, therefore the results of both sexes were pooled together. In all cases mice were acclimatized to the behavioral room for at least 20 min before testing and all apparatus were cleaned with 70% ethanol between animals. Behavioral evaluation started 2 weeks after the first tamoxifen injection or 2 weeks after AAV2.Cre virus delivery into the mHb. The different behavioral tests were performed in the following order with 4 days between each test: open field test, elevated-plus maze, innate fear response, passive avoidance. In a different set of mice, the following behavioral tests were performed with 7 days between each test: open field test, context fear conditioning, cued fear conditioning.

#### Open field test

Mice were placed in the center of a 48cm x 48cm transparent acrylic box equipped with lightbeam strips (TSE Actimot system) and allowed to move freely for 10 min. General locomotor parameters including distance travelled, time moving and distance and time spent in the center of the arena were calculated using Actimot Software.

#### Elevated-plus maze

The apparatus consisted of a cross-shaped maze with two open arms opposite to two arms enclosed by lateral walls (70 cm x 6 cm x 40 cm) elevated 70 cm above the floor. Each animal was placed in the central square (6 cm x 6 cm) facing one of the open arms away from the experimenter and allowed to explore the maze for 5 min. The movement of the animals was recorded using the Ethovision XT-10 tracking software and the number of entries and the staying time in each of the arms were analyzed.

#### Innate fear measurement

The innate freezing response of the animals after a foot-shock was evaluated using a TSE multiconditioning system (TSE Systems). Each animal was placed in a squared behavioral arena made of plexiglass, and after 3 minutes of habituation the mouse received 3 consecutive footshocks separated by 1 min intervals (0.7 mA; 1s). Animals were left in the arena for 2 extra minutes and the percentage of freezing behavior during each minute interval was analyzed.

#### Passive avoidance

The apparatus chamber used in this test was composed by two compartments (a dark and an illuminated one) connected through an automatic door (TSE multiconditioning system, TSE Systems). On the acquisition/conditioning day animals were place on the illuminated compartment and 30 s after the door was opened. When the animals crossed to the dark chamber, the door was closed and they received a mild foot-shock (1s, 0.7 mA). The day after the mice were placed again on the illuminated compartment, 30 s later the door was opened and the latency to enter the dark side was measured with a maximum latency time of 600s.

#### Context Fear conditioning

Mice were exposed to context A (square box with one transparent side and three black sides) for 3 min ending with a foot-shock (2s, 0.7mA). The day after, the animals were exposed to the same context A for 3 min and the freezing was measured using the automatic counting system in the TSE system.

#### Cued Fear conditioning

Mice were exposed to context A (square box with one transparent side and three black sides) for 2 min and then a tone was played for 20s ending with a foot-shock (0.7mA, 2s). The day after the animals were placed in context B (a transparent cylinder box) and after 1 min they were exposed to the same tone for 5 min. The freezing was measured using the automatic counting system in the TSE system.

### Statistical analysis

Data are expressed as mean and standard errors (s.e.m). No statistical methods were used to predetermine sample sizes but our sample sizes are similar to those generally used in the field. Following normality test and homogeneity variance (F-test or Kolmogorov-Smirnov test with Dallal-Wilkinson-Lilliefor p value), group comparison was made using a unpaired student t-test, one-way or two-way ANOVA as appropriate followed by Bonferroni or Tukey’s multiple comparison test for normally distributed data. Mann–Whitney U test or Kruskal-Wallis followed by Dunn’s multiple comparison test was used on non-normal distributed data. Differences were considered significant for p< 0.05.

## Supporting information

Main text

## Acknowledgements

The authors would like to thank Maria Christina Sergaki and Wei Wang for technical assistance during the early stages of the project; Mart Saarma and Jaan-Olle Andressoo (University of Helsinki, Finland) for *Gfra1*^fx/fx^ mice; Linda Thors, Emma Wallet and personnel from the KMB animal facility of Karolinska Institutet for their help with animal care; IIna Eleonoora Korkala for DAPI cell counts; and all CIB Lab members for comments and suggestions. Support for this research was provided by grants to C.F.I. from the Swedish Research Council (2016-01538 and 2020-01923) and the National Research Foundation of Singapore (R-711-000-052-281).

## Author contributions

D.F.S. performed the majority of the experimental work; F.K. performed the PLA experiments; M.D. and K.P. performed the optogenetic experiments; A.A. provided technical support for mouse genotyping as well as behavioral and histology studies; L.K. contributed to initial signaling studies; D.F.S. and C.F.I. designed the experiments and wrote the paper.

## Abbreviations

BAC: Bed nucleus of the anterior commisure
ChAT: Choline acetyltransferase
CNO: Clozapine-N-oxide
CR: Calretinin
DREADD: Designer Receptor Exclusively Activated by Designer Drug
dmHb: Dorsal medial habenula
EPSCs: Excitatory post-synaptic currents
FR: Fasciculus retroflexus
GDNF: Glial cell derived neurotrophic factor
GFRα1: Glial cell derived neurotrophic factor receptor alpha 1
IPN: Onterpeduncular nucleus
LHb: Lateral Habenula
mHb: Medial habenula
NCAM: Neural cell adhesión molecule
PLA: Proximity ligation assay
SNpc: Substantia nigra pars compacta
SP: Substance P
TS: Triangular septum
VGAT: Vesicular GABA transporter
VGlut1: Vesicular glutamate transporter 1
VGlut2: Vesicular glutamate transporter 2
vmHb: ventral medial habenula
VTA: Ventral tegmental área

