## Supplementary material for "Adult medial habenula neurons require GDNF receptor GFRα1 for synaptic stability and function": Main text

### Figure S1

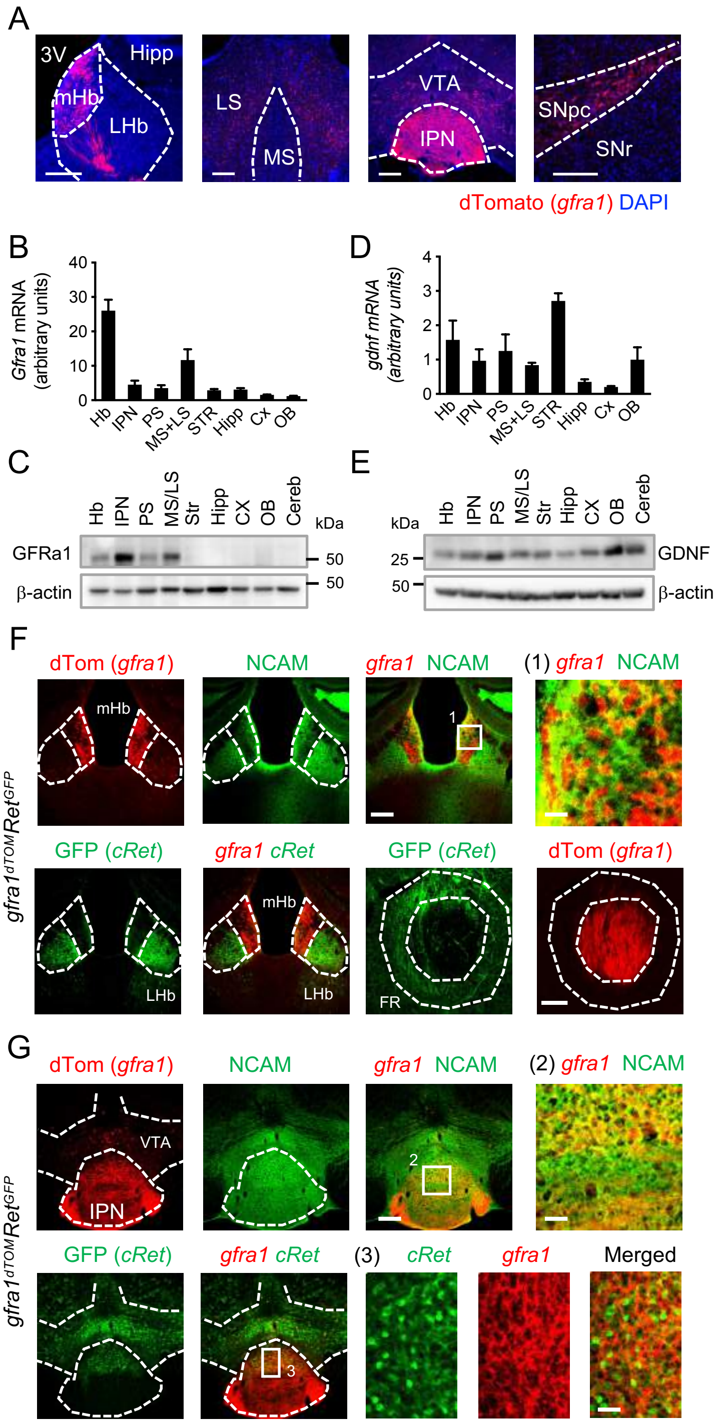


**Figure S1. Characterization of GFRα1 and co-receptor expression in the adult mouse brain.**
**A)** dTomato epifluorescence (red) in coronal sections of *Gfra1$^{dTOM}$* mouse brain injected with tamoxifen at 3 months counterstained with DAPI (blue). Hipp: hippocampus; IPN: interpeduncular nucleus; MS: medial septum; mHb: medial habenula; LHb: lateral habenula; LS: lateral septum; SNpc: substantia nigra *pars compacta*; SNr: substantia nigra *reticulata*. Scale bars, 200μm.
**B,D)** Expression levels of *gfra1* (B) and *gdnf* (D) mRNAs quantified by qPCR and normalized against *18S* in brain areas dissected from 3 month old C57BL6/J mice (n=5) . Cx: cortex; Hb: habenula; OB: olfactory bulb; PS: posterior septum; STR: striatum.
**C,E)** Immunoblots of whole protein extracts of 3 month old C57BL6/J mice probed for GFRα1 (C) and GDNF (E). β-actin was probed as loading control.
**F,G)** dTomato epifluorescence (red), GFP (green) and NCAM (green) immunolabelling in coronal sections of the mHb and FR (F) and IPN (G) of *gfra1$^{dTOM}$Ret$^{GFP}$* mouse injected with tamoxifen at 3 months. Scale bars, 200μm (mHb) and 30μm (inset 1), 50 μm (FR), 200μm (IPN) and 40μm (insets 2 and 3).

### Figure S2

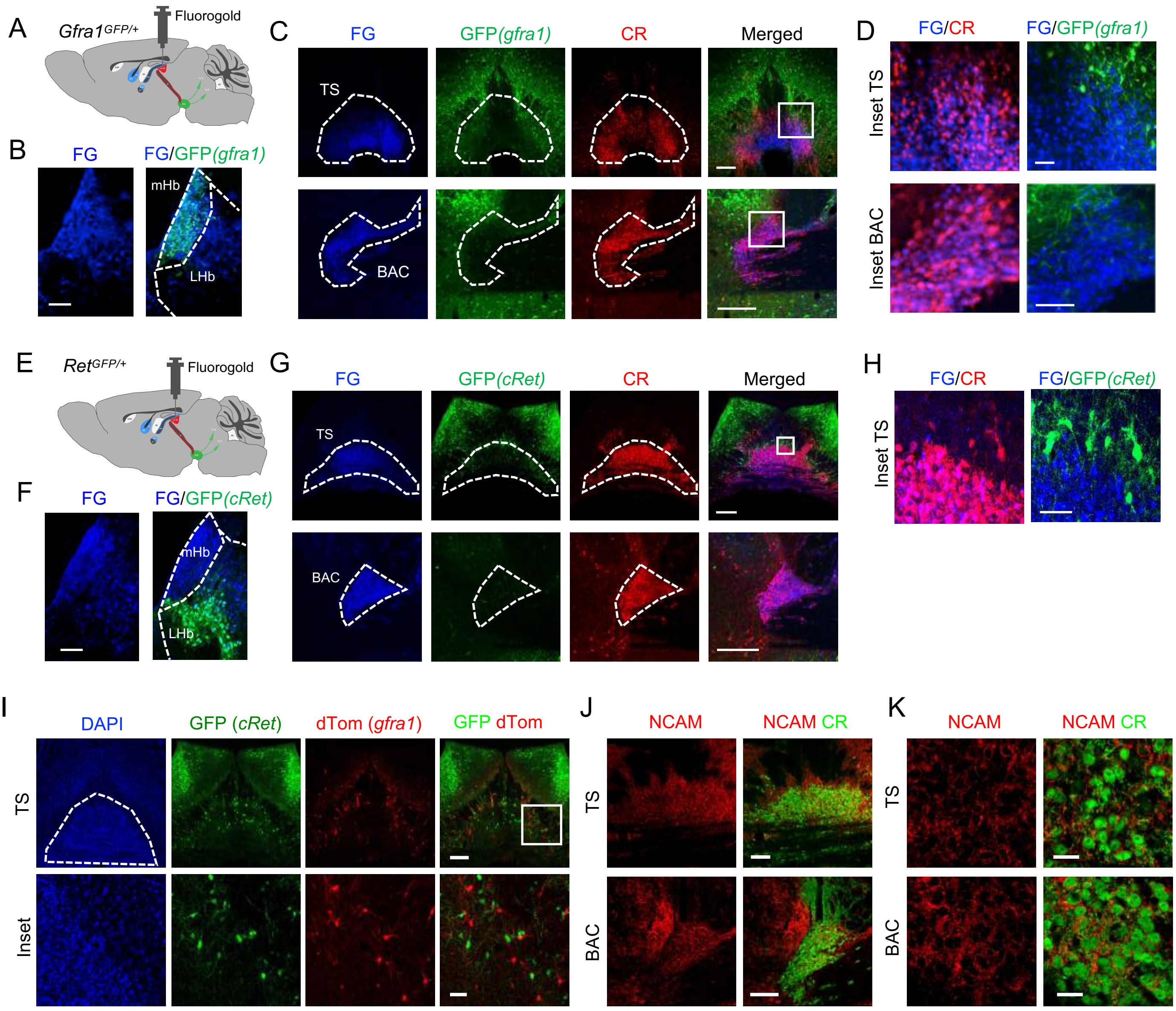


**Figure S2**. **Expression of GFRα1 and co-receptors in the TS and BAC.**
**A,E)** Schematic representation of a sagittal brain section of a 3 month old *Gfra1*$^{GFP}$ (A) or *Ret*$^{GFP}$ (E) mouse injected with fluorogold (FG) in the mHb.
**B,F)** GFP (green) immunolabelling in coronal sections of the mHb of a *Gfra1*$^{GFP}$ (B) or a *Ret*$^{GFP}$ (F) mouse injected with FG (blue) in the mHb. Scale bars, 100μm.
**C,D,G,H)** Calretinin (CR, red) and GFP (green) immunolabelling in coronal sections of the TS and BAC from a *Gfra1*$^{GFP}$ (C,D) or a *Ret*$^{GFP}$ (G,H) mouse injected with FG (blue) in the mHb. Scale bars, 200μm (C,G) and 50μm (D,H).
**I)** dTomato epifluorescence (red) and GFP (green) immunolabelling in a coronal section counterstained with DAPI (blue) of the TS of a *Gfra1*$^{dTOM}$*Ret*$^{GFP}$ mouse treated with tamoxifen at 3 months. Scale bars, 200μm and 50μm (inset).
**J,K)** NCAM (red) and CR (green) immunolabelling in coronal sections of the TS and BAC of a 3 month old C57BL6/J mouse. Scale bars, 100μm (J) and 20μm (K).

### Figure S3

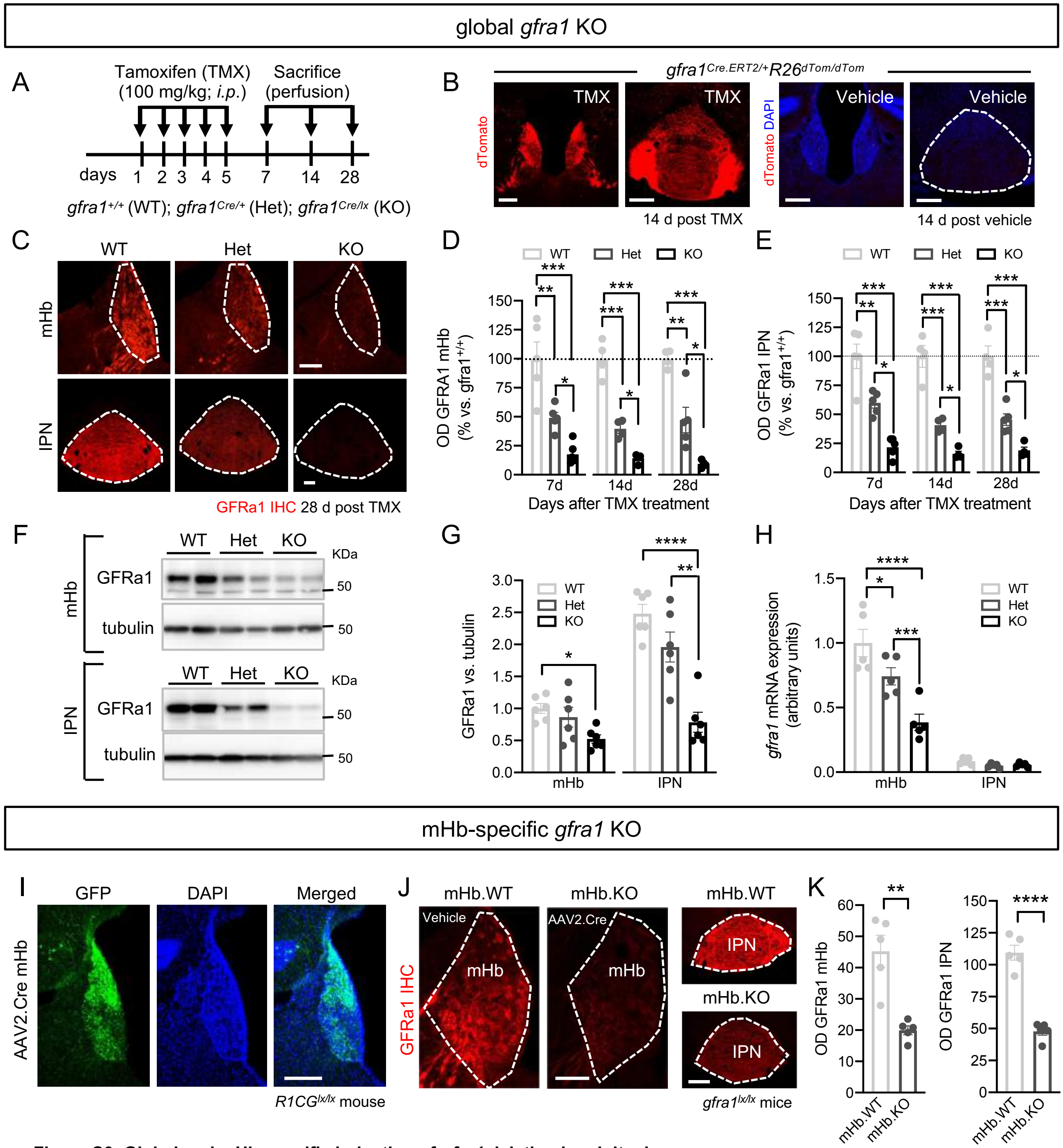


**Figure S3**. **Global and mHb-specific induction of *gfra1* deletion in adult mice.**
**A)** Schematics of the global *gfra1* KO mouse model. **B)** dTomato epifluorescence (red) in coronal sections counterstained with DAPI (blue) of the mHb and the IPN of *gfra1*$^{dTOM}$ mouse treated with tamoxifen or vehicle at 3 months. Scale bars, 200µm. **C)** Immunolabelling for GFRα1 (red) in coronal sections of the mHb and the IPN of WT, Het and KO mice. Scale bars, 100µm. **D,E)** Quantification (±SEM) of GFRα1 optical density (OD) in the mHb (D) and the IPN (E) of WT, Het and KO mice normalized against the WT group. N= 4-6 animals/group; 1-way ANOVA analysis followed by Tukey's post-hoc test at each time point; * $p< 0.05$; ** $p< 0.01$; *** $p< 0.001$. **F,G)** Immunoblots of total protein fraction from mHb and IPN of WT, Het and KO mice probed for GFRα1 (F) and αtubulin. Quantifications (±SEM) of GFRα1 levels were corrected for αtubulin levels and normalized to levels in WT samples. N=6 mice per group; 1-way ANOVA followed by Tukey's post-hoc test; * $p< 0.05$; ** $p< 0.01$; **** $p< 0.0001$. **H)** Quantification by qPCR of *Gfra1* mRNA expression levels (±SEM) in the mHb and the IPN of WT, Het and KO mice. *Gfra1* levels were corrected for *18S* levels and normalized to levels in the mHb of WT samples. N=5 mice per group, 2-way ANOVA followed by Tukey's post hoc test; * $p< 0.05$; *** $p< 0.001$; **** $p< 0.0001$. **I)** GFP (green) immunolabelling in coronal sections counterstained with DAPI (blue) of the mHb of a *R1CG*$^{lx/lx}$ mouse injected with an AAV.CMV.Cre virus in the mHb. Scale bar, 100µm. **J)** GFRα1 immunolabelling (red) in coronal sections of the mHb and the IPN of *gfra1*$^{lx/lx}$ mice injected with vehicle (mHb.WT) or AAV.Cre (mHb.KO) in the mHb. Scale bars, 50µm (mHb) and 200µm (IPN). **K)** Quantification (±SEM) of GFRα1 OD in the mHb and the IPN of mHb.WT and mHb.KO mice. N=5 animals per group; Student's t-test; ** $p< 0.01$; **** $p< 0.0001$.

### Figure S4

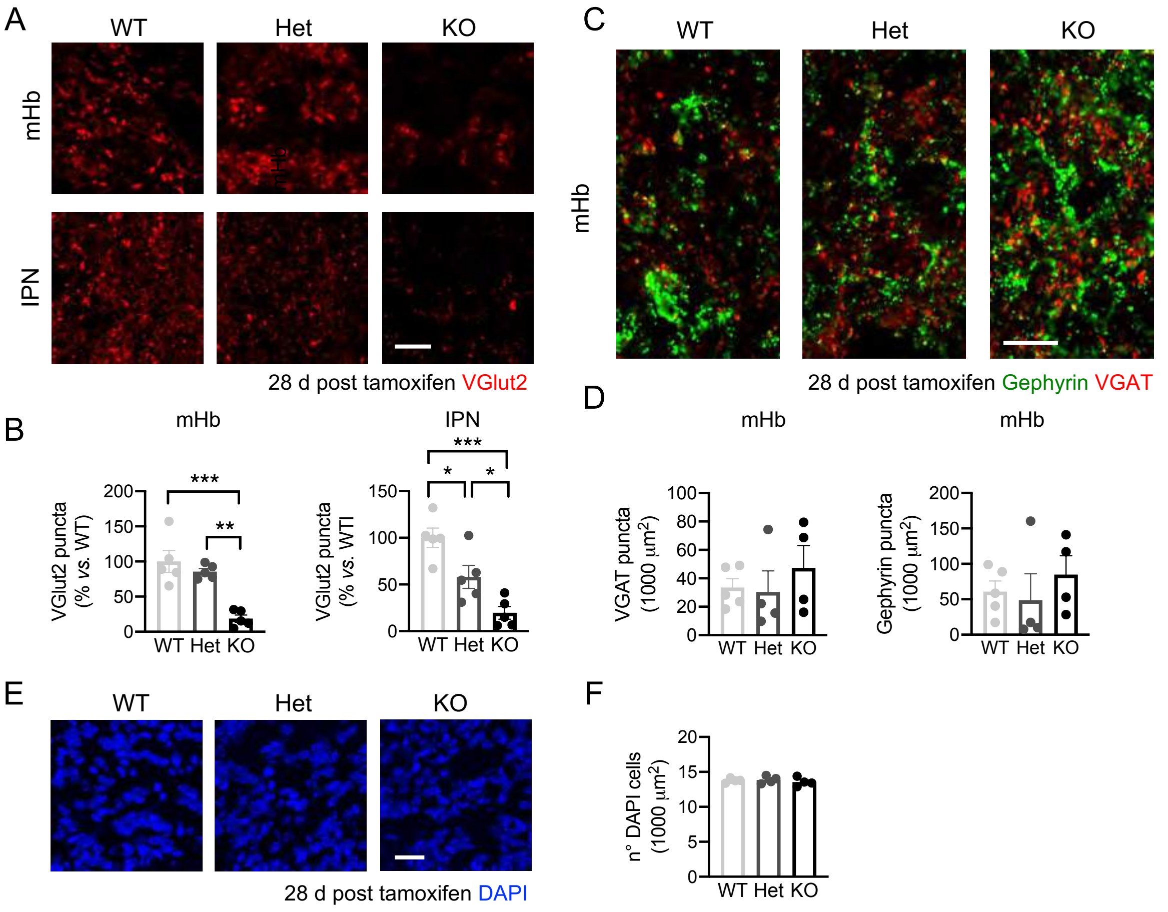


**Figure S4. Quantification of VGlut2, VGAT and gephyrin puncta and number of DAPI cells after global ablation of GFRα1.**

**A)** VGlut2 (red) immunostaining in coronal sections of the mHb and the IPN of WT, Het and KO mice. Scale bar, 10µm.

**B)** Quantification (±SEM) of puncta with immunoreactivity for VGlut2 in mHb and IPN. N=5 mice per group (25-30 images per mouse); 1-way ANOVA followed by Tukey's post hoc test; * $p < 0.05$, ** $p < 0.01$, *** $p < 0.001$.

**C)** VGAT (red) and gephyrin (green) immunostaining in coronal sections of the mHb of WT, Het and KO mice. Scale bar, 10µm.

**D)** Quantification (±SEM) of puncta with immunoreactivity for VGAT and gephyrin in the mHb. N=4-5 mice per group (25-30 images per mouse).

**E)** DAPI (blue) counterstaining in coronal sections of the mHb of WT, Het and KO mice. Scale bar, 20µm.

**F)** Quantification (±SEM) of number of DAPI cells in the mHb. N=4 mice per group (32 images per mouse).

### Figure S5

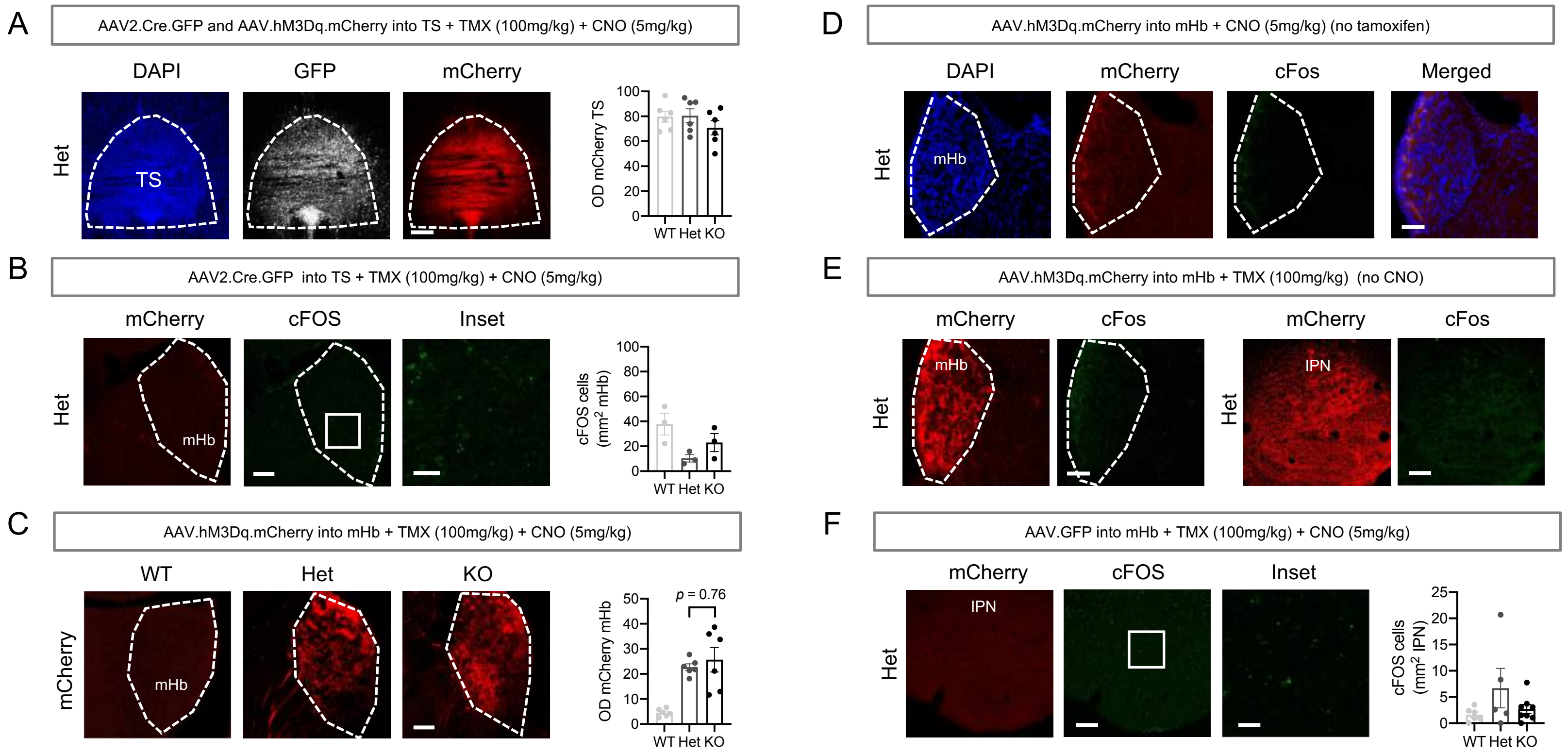


**Figure S5. Control groups of the chemogenetic experiments.**
Mice were injected with viral vectors in the mHb or the TS and treated with tamoxifen, CNO or their vehicles as indicated in each panel.

**A)** mCherry epifluorescence (red), GFP (grey) immunolabelling and counterstaining in DAPI (blue) in a coronal section of the TS of a Het mouse (scale bar, 200µm). Graph shows the quantification (±SEM) of the optical density (OD) for mCherry in the TS of WT, Het and KO mice. N= 6 mice per group (4-6 images per mouse).

**B)** mCherry epifluorescence (red) and GFP immunolabelling (green) in a coronal section of the mHb of a Het mouse. Scale bars, 100µm (mCherry) and 30µm (inset). Graph shows quantification of cFOS positive cells (±SEM) in WT, Het and KO mice. N=3 mice per group (14-16 images per mouse).

**C)** mCherry epifluorescence (red) in coronal sections of the mHb of WT, Het and KO mice. Scale bar, 100µm. Graph shows quantification (±SEM) of OD for mCherry in the mHb. N=6 mice per group (14-16 images per mouse); 1-way ANOVA followed by Tukey's post-hoc test.

**D)** mCherry epifluorescence (red) and cFOS immunostaining (green) in a coronal section counterstained with DAPI (blue) of the mHb of a Het mouse. Scale bar, 100µm.

**E)** mCherry epifluorescence (red) and cFOS immunostaining (green) in coronal sections of the mHb and the IPN of a Het mouse. Scale bars, 200µm and 50µm (inset).

**F)** mCherry epifluorescence (red) and cFOS immunostaining (green) in a coronal section of the IPN of a Het mouse. Scale bars, 100µm (mHb) and 200µm (IPN). Graph shows quantification (±SEM) of cFOS cells in the IPN. N=5-8 mice per group (4-6 images per mouse).

### Figure S6

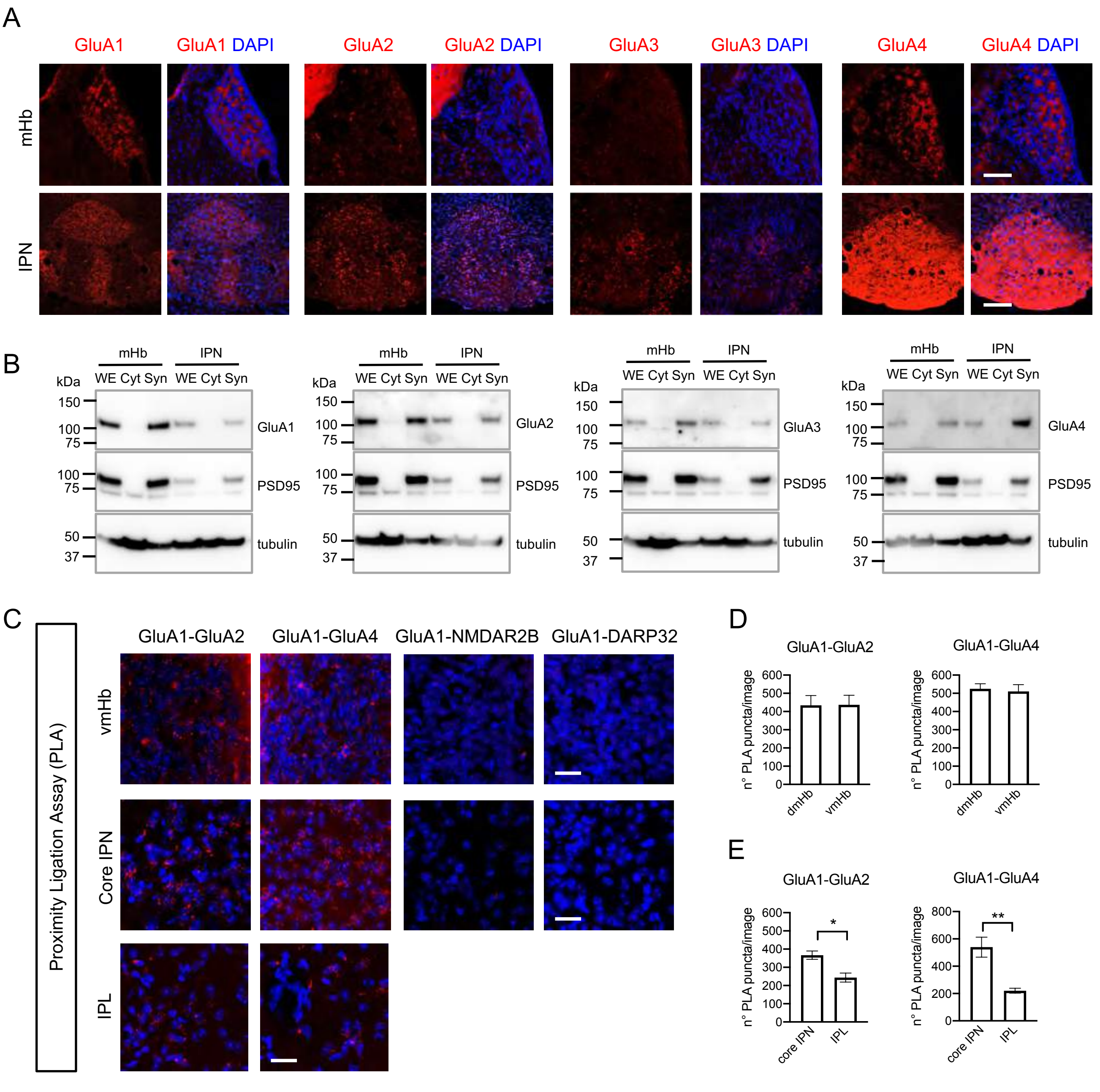


**Figure S6. Characterization of GluA1-4 expression in the mHb and IPN of C57BL6/J adult mice.**

**A)** GluA1-4 immunostaining (red) and counterstaining in DAPI (blue) in coronal sections of the mHb and the IPN of a 3 month old C57BL6/J mouse. Scale bars, 150μm (mHb) and 300μm (IPN).

**B)** Immunoblots of whole (WE), cytosolic (Cyt) and synaptosome (Syn) protein extracts from the mHb and IPN of 3 month old C57BL6/J mice probed against GluA1-4. PSD95 and tubulin were probed as loading controls.

**C)** Proximity Ligation Assay (PLA) signals (red) for GluA1-GluA2 and GluA1-GluA4 complexes in coronal sections of mHb and IPN from 3 month old C57BL6/J mice. Counter-staining with DAPI appears in blue. NMDAR2B and DARP32 were used as negative controls. Scale bars, 20 μm.

**D,E)** Quantification (±SEM) of PLA puncta for GluA1-GluA2 and GluA1-GluA4 complexes in the dorsal and ventral mHb (D) and the lateral and core IPN (E) from 3 month old C57BL6/J mice. N=4 mice (16 images per mHb and 18 images per IPN). Student's T-test; * $p<0.05$; ** $p<0.01$.

### Figure S7

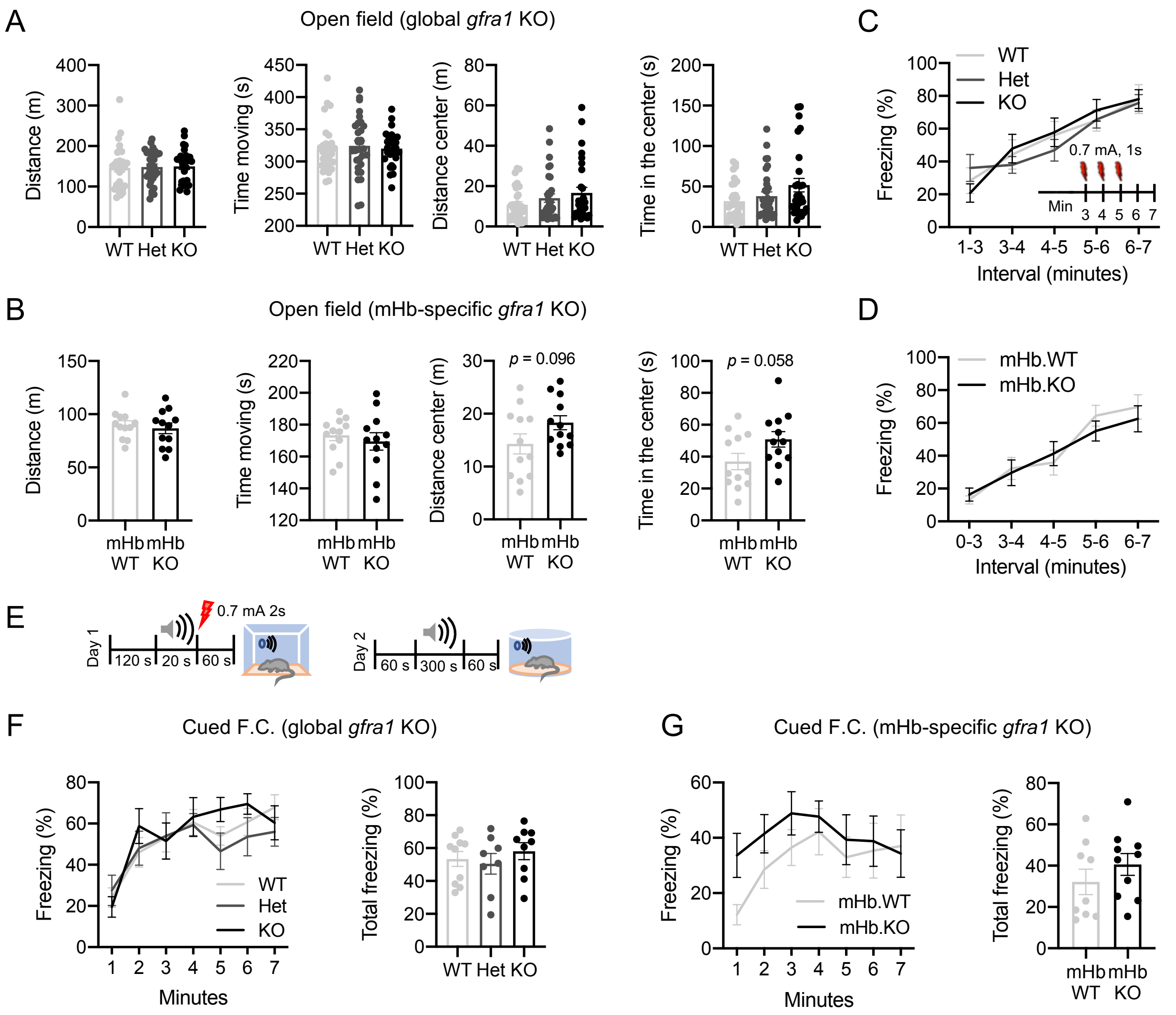


**Figure S7. Acute loss of gfra1 in adult mHb neurons does not alter the behavioral responses in the open field, the innate freezing response or a cued fear conditioning paradigm.**

**A,B)** Total distance, time moving and distance and time spent in the center of the open field arena (±SEM) for WT, Het and KO mice (A) and mHb.WT and mHb.KO mice (B). N=28-31 mice per group, Kruskal-Wallis test followed by Dunn's multiple comparison test (A). N=12 mice per group, Student's t-test (B).

**C,D)** Freezing responses (±SEM) after 3 consecutive foot-shocks for WT, Het and KO mice (C) and mHb.WT and mHb.KO mice (D). N=10-13 mice per group, 2-way ANOVA followed by Bonferroni post-hoc test (C). N=12 mice per group, Student's t-test.

**E,G**) Schematic (E), freezing time-course and total freezing time (±SEM) for WT, Het and KO mice (F) and mHb.WT and mHb.KO mice (G) in the Cued Fear Conditioning test. N=8-10 mice per group, 2-way ANOVA and 1-way ANOVA (F). N=9-10 mice per group, 2-way ANOVA and Student's t-test (G).
